# *MYC–NFATC2 axis* maintains cell cycle and mitochondrial function in AML cells

**DOI:** 10.1101/2023.11.15.567209

**Authors:** S.D. Patterson, M.E. Massett, X. Huang, H.G. Jørgensen, A.M. Michie

## Abstract

Acute myeloid leukaemia (AML) is a clonal haematological malignancy affecting the myeloid lineage with generally poor patient outcomes, owing to the lack of targeted therapies. The histone lysine demethylase 4A (KDM4A) has been established as a novel therapeutic target in AML, due to its selective oncogenic role within leukaemic cells.

We identify that the transcription factor *NFATC2* is a novel binding and transcriptional target of KDM4A in the human AML THP-1 cell line. Further, cytogenetically diverse AML cell lines, including THP-1, were dependent on *NFATC2* for colony formation *in vitro*, highlighting a putative novel mechanism of AML oncogenesis.

Our study demonstrates that *NFATC2* maintenance of cell cycle progression in human AML cells was driven primarily by *CCND1*. Through RNA-seq and ChIP-seq, *NFATC2* was shown to bind to the promoter region of genes involved in oxidative phosphorylation and subsequently regulate their gene expression in THP-1 cells. Furthermore, our data show that *NFATC2* shares transcriptional targets with the transcription factor c-MYC, with *MYC* knockdown phenocopying *NFATC2* knockdown. These data suggest a novel co-ordinated role for *NFATC2* and *MYC* in the maintenance of THP-1 cell function, indicative of a potential means of therapeutic targeting in human AML.

**Graphical Abstract:** 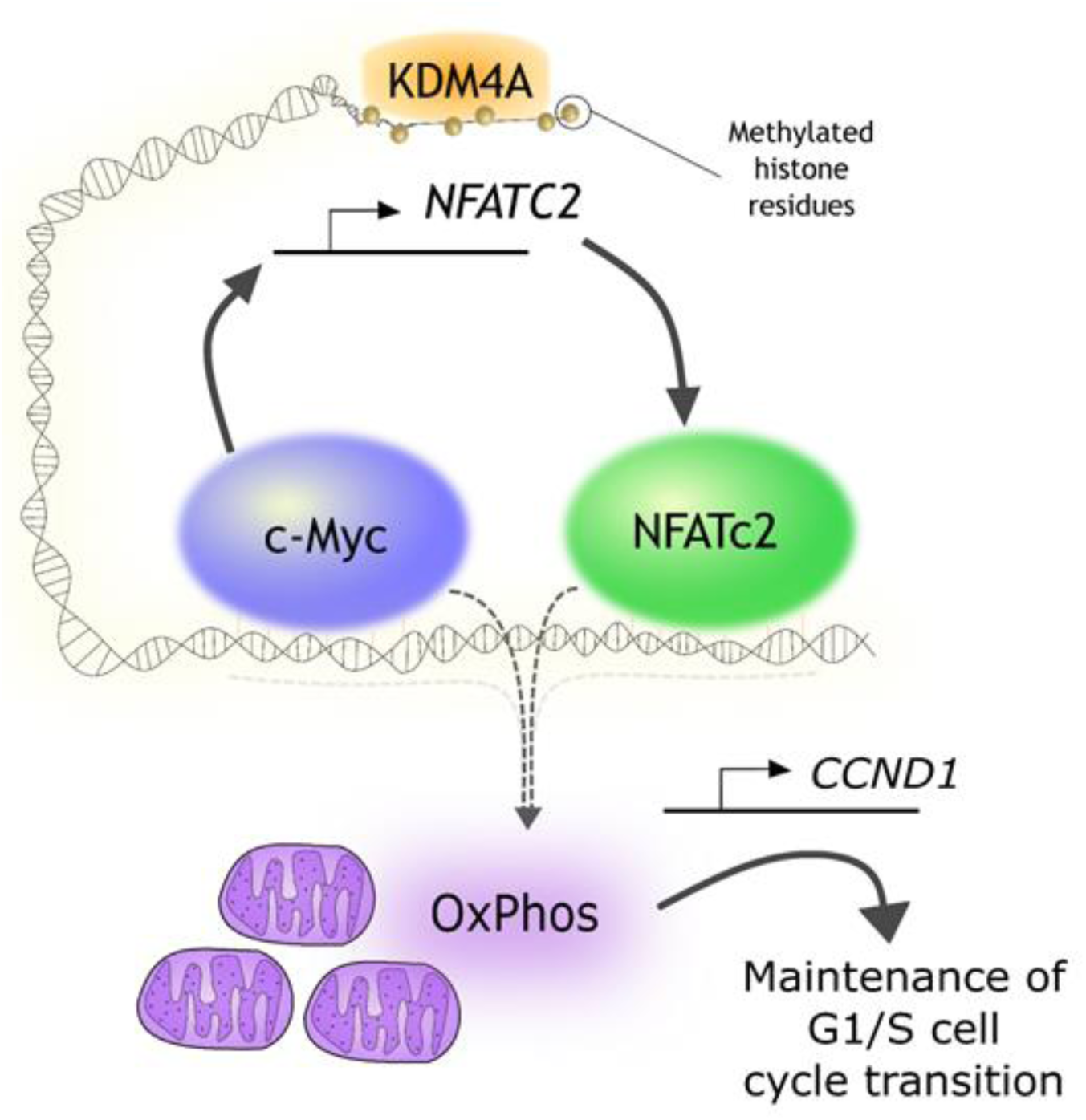

Acute myeloid leukaemia (AML) cells of diverse cytogenetic backgrounds are dependent on *NFATC2* for survival; in THP-1 cells, *NFATC2* is downstream of the epigenetic regulator *KDM4A*. *NFATC2* promotes G1/S phase transition in the cell cycle and oxidative phosphorylation. In addition, *NFATC2* functions downstream of transcription factor *MYC*, and maintains *CCND1* expression.

## Introduction

Acute myeloid leukaemia (AML) is a haemopoietic malignancy of clonal origin with poor outcomes; 5-year survival is as low as 13% [1], and while most AML patients are over 60 years of age, there is incidence of AML at all ages [2]. Despite improvements to patient outcomes over recent years, most of these have been observed in patients under 60, due to generally poorer tolerance of chemotherapy in older patients [3] and the significant cytogenetic heterogeneity between patients’ disease, the latter of which making it challenging to develop effective targeted therapies. As such, novel targeted therapies which can successfully eliminate pathogenic AML cells in stratified groups of patients are necessary.

Epigenetic modifiers have been characterised as having key roles in the pathogenesis of AML [4]. The histone lysine demethylase 4A (KDM4A), which regulates transcriptional activity via demethylation of histone 3 lysine residues, including 9, 27 and 36, has been characterised as a ‘master’ regulator of AML cell survival mechanisms. Massett *et al*. demonstrated that genetic depletion of KDM4A or small molecule inhibition of KDM4 family proteins was detrimental to AML cell survival, while sparing healthy/normal haemopoietic stem cells (HSCs) [5]. Supporting this, other studies showed that small molecule pan-inhibition of KDM4 and KDM6 family proteins was detrimental to cell survival [6]. Furthermore, another member of the KDM family, KDM1A, was shown to be essential for the survival of AML cells driven by MLL-AF9, a leukaemogenic gene fusion product, through maintenance of a pro-oncogenic transcriptional signature [7]. There is also value in exploring the transcriptional mechanisms downstream of KDM4A, to understand its mechanisms of oncogenesis and identify novel targetable leukaemogenic pathways, which may support the development of novel precision medicine approaches; for example, for stratifying patients that have differing responses to KDM4A inhibitors or demethylase-targeting therapies.

The Nuclear Factor of Activated T cells (NFAT) family consists of five members, in which NFATc1-4 proteins function downstream of calcium signalling, while NFAT5 is responsive to osmotic stress. Physiologically, inactive NFAT proteins reside in the cytoplasm in a heavily phosphorylated state, and activation of calcium-coupled surface receptors triggers a signalling cascade via phospholipase C (PLC), which promotes calcium influx and subsequent dephosphorylation of NFAT, primarily by the phosphatase calcineurin. Dephosphorylated NFATs then translocate to the nucleus, where they bind DNA as a monomer and activate transcription; the most well-characterised example of this being at the IL-2 promoter in T cells [8]. However, their role in myeloid cells has received limited attention. NFAT expression is downregulated during the differentiation of CD34^+^ HSCs towards mature myeloid cells, and different NFAT family members have been found to have divergent roles in cell fate [9, 10] and regulation of the cell cycle [11, 12]. There is also growing evidence that NFAT signalling can enable therapy resistance in myeloid leukaemias [13, 14] and enable AML development, such as in AML cells with the internal tandem duplication in FLT3 (FLT3^ITD^) [15]. In this study, we identified the *NFATC2* promoter as a novel binding and transcriptional target of KDM4A. We used genetic knockdown (KD) and global sequencing approaches in human AML cell lines, including a number of cytogenic AML subtypes, to characterise novel mechanisms of *NFATC2*-regulated oncogenic maintenance in AML, indicative of a putative targetable mechanism for further study.

## Material and Methods

### Cell culture

The suspension cell lines THP-1, HL-60, MV4-11, MOLM-13, Kasumi-1 (all ATCC; RRIDs: CVCL_0006, CVCL_0002, CVCL_0064, CVCL_2119, CVCL_0589, respectively) and OCI-AML3 (DSMZ; RRID: CVCL_1844) were cultured in HEPA-filtered incubators at 37°C and with 5% CO_2_, in RPMI supplemented with 2 mM L-glutamine and 10-20% (V/V) FBS, as recommended by the supplier. The adherent cell lines HEK-293T (ATCC; RRID: CVCL_0063) and HEK-293T/phoenix-AMPHO (ATCC; RRID: CVCL_H716) were maintained in DMEM with 2 mM L-glutamine and 10% FBS, and sub-cultured using Trypsin-EDTA (0.25%; Thermo Fisher Scientific). HEK-293T/phoenix-AMPHO cells were selected for stable expression of retroviral vectors using Hygromycin B (200 µg/mL) and Diphtheria toxin (2 µg/mL; Merck) [16]. All cell lines were authenticated in the previous three years by monitoring growth curves and cell morphology; further information is provided below. All cultures were regularly validated to be mycoplasma-negative.

### Virus generation

Lentiviral supernatants were generated by transfection of HEK-293T cells in 6-well plates using PEI-mediated transfection. In brief, for lentivirus generation, PEI was complexed with plasmids in a total mass ratio of 12 µg PEI: 4 µg DNA/well in serum-free DMEM (1.8 µg shRNA (pLKO.1):1.8 µg packaging (psPAX2):0.4 µg envelope (pMD2.G) plasmids). These were incubated for 30 min at RT, before adding dropwise to each well, which contained HEK-293T culture media. Media were changed after 24 hr and lentiviral supernatants were harvested subsequently at two 24 hr intervals and filtered using 0.45 µm filters. Retroviral supernatants were generated in the same manner, using HEK-293T/phoenix-AMPHO cells, with 1.8 µg shRNA (pLKO.1)/well.

Cell lines were transduced with lenti/retrovirus by mixing cell suspension (density 1-2×10^5^/mL, depending on manufacturer-suggested minimum seeding density) with lenti/retroviral supernatant in a 1:1 volumetric ratio and 8 µg/mL hexadimethrine bromide (Merck), before centrifugation at 900g for 30 min and returning to incubation at 37°C with 5% CO_2_. At 24 and 48 hr post-transduction, lentivirus-transduced cell suspensions were selected using puromycin (concentration determined empirically per cell line; Merck). Experimental timepoints are referred to as “post-selection”, relative to the first addition of puromycin (0 hr). Retrovirus-transduced cells were selected at 48 hr post-transduction for positive GFP expression using fluorescence-activated cell sorting.

### RNA-seq

For RNA-sequencing (RNA-seq), the cDNA libraries were prepared from prepared RNA by Novogene using a Next® Ultra^TM^ RNA Library Preparation Kit (NEB) and sequenced using a NovaSeq 6000 platform (Illumina). Reads were aligned to the hg38 human genome build using STAR and expression was quantified using the HTseq package; reads are referred to in the units Fragments Per Kilobase of transcript per Million mapped reads (FPKM). Differential expression between samples transduced with sh*KDM4A* or sh*NFATC2* and the NTC was conducted in a pairwise manner, using the DESeq2 R package, which utilised 3 biological replicates in the calculation. Next, parsing and filtering of differential expression data, and production of most diagrams were conducted in Rstudio. Further detail of analyses is included in the Supplementary Methods section.

### Chromatin Immunoprecipitation coupled with sequencing (ChIP-seq)

*NFATC2*-bound DNA was precipitated from untreated THP-1 cells (11×10^6^ cells/IP). Cells were fixed using formaldehyde (1% V/V in H_2_O) for 10 min at RT before supplementation with 125 mM glycine solution (Merck) and agitation for 5 min. Cytosolic lysate was generated from fixed cells using cytoplasmic lysis buffer (10 mM Tris pH 8.0/10 mM NaCl/0.4% (V/V) IGEPAL CA-630/1X protease inhibitor cocktail/10 µM DIFP; Merck) and removed, before generation of nuclear lysate using nuclear lysis buffer (50 mM Tris pH 8.0/EDTA 50 mM/SDS 0.8% (W/V)/1X protease inhibitor cocktail/10 µM DIFP; Merck), which was then sonicated using an EpiShear^TM^ Probe Sonicator (Active Motif), with 18 bursts at 30% amplitude for 30 sec per burst. Sheared lysates were pre-cleared using 2 µg rabbit IgG antibody (Thermo Fisher Scientific) per sample for 1 hr at 4°C, before removing IgG-bound lysate by incubating with protein G magnetic Dynabeads® (Thermo Fisher Scientific) for 1 hr and then removal by incubation with a magnetic stand. Next, 2 µg antibody targeting either NFATc2 or rabbit IgG was added to pre-cleared supernatants and incubated overnight at 4°C.

Following this, bead-bound DNA was magnetically isolated, washed and then eluted using an Ipure V2 Kit, as per protocol, and sequencing libraries were prepared using a MicroPlex Library Preparation Kit (both Diagenode). The libraries were purified using AMPure® XP beads (Beckman Coulter) and sequenced by Novogene using a NovaSeq 6000 platform (Illumina).

Raw data were processed using tools in the Galaxy platform and subsequent analyses performed in RStudio (version 4.3). Further details of analyses are included in the Supplementary Methods.

### Colony-forming cell assay

For each condition, 1000 cells were seeded in MethoCult^TM^ H4230 semi-solid media, supplemented with IMDM to 20% (V/V) in technical triplicate, and a minimum of three independent biological replicates, and lentivirus-transduced cells were seeded with puromycin also. Assays were then incubated at 37°C with 5% CO_2_. Colonies were counted and imaged at 7-10 days post-seeding using an EVOS cell imaging system.

### Metabolic assays: MitoSOX, CellTiter-Glo, Alamar Blue

For the MitoSOX assay, cells were incubated with 1 mL 500 nM MitoSOX red (Thermo Fisher Scientific) in HBSS for 30 min at 37°C and with 5% CO_2_, before washing in HBSS and staining with DAPI (1 µg/mL). Using flow cytometry, live (DAPI^-^) and GFP^+^ cells were subset and gated for MitoSOX positivity.

To assess ATP content, cell suspensions (plated at equivalent cell densities) were incubated with an equivalent volume of CellTiter-Glo (Promega) working reagent, in technical triplicate, in an opaque 96-well plate for 10 min. The luminescent signal was measured using a SpectraMax® M5 Plate Reader.

For the Alamar Blue assay, Alamar Blue dye (Merck) was incubated with cell suspensions (plated at equivalent cell densities) at 50 µM, in technical triplicate, at 37°C with 5% CO_2_ for 4 hr. The absorbance was measured at excitation 535 nm and emission 590 nm using a SpectraMax® M5 Plate Reader.

### Statistical Analyses and Software

For the statistical testing of differences between multiple groups, a one-way ANOVA test with post-hoc Dunnett’s tests for a comparison of means between the treatment group and control was used. A two-way ANOVA with post-hoc Dunnett’s tests for a comparison of means between the treatment group and control was used for inter-group comparisons with multiple factors. For multiple comparisons at differing timepoints, a repeated measures ANOVA test was used. In the case of comparing only two groups, an unpaired, two-sided t-test was used. p<0.05 was considered significant. For the analyses of RNA-seq and ChIP-seq data, the tests are specific to the package used and are stated in the figure legends. In each case, appropriate correction for multiple hypothesis testing was applied by the package used.

Graphs were assembled and the associated statistical tests were conducted using GraphPad Prism software (version 10.0). RNA-seq and ChIP-seq data, in addition to the patient datasets, were analysed using a combination of the Galaxy platform and Rstudio (version 4.3).

All additional standard methods are detailed in the Supplementary File.

## Results

### *NFATC2* is a downstream transcriptional target of KDM4A in THP-1 cells

We have previously demonstrated that human AML cells, including THP-1 cells, are dependent on *KDM4A* for survival [5]. RNA-seq was performed in THP-1 cells with *KDM4A* KD to identify the pathways that were deregulated by depletion of KDM4A. Analysis of this dataset (GSE125375) using SPIA, a signalling pathway enrichment tool, yielded 137 pathways, 32 of which, relating to intracellular signalling, were included for downstream analyses, to focus specifically on the elucidation of intracellular signalling perturbations; the topmost deregulated pathways are shown in Figure 1A. The ‘Wnt signaling pathway’ was the most inhibited pathway after sh*KDM4A* transduction, according to the descriptors of SPIA (total accumulated perturbation of the pathway (tA) = -7.457; false discovery rate for pathway perturbation (pGFdr) <0.001), compared to the non-targeting control (NTC) shRNA.

**Figure 1.**
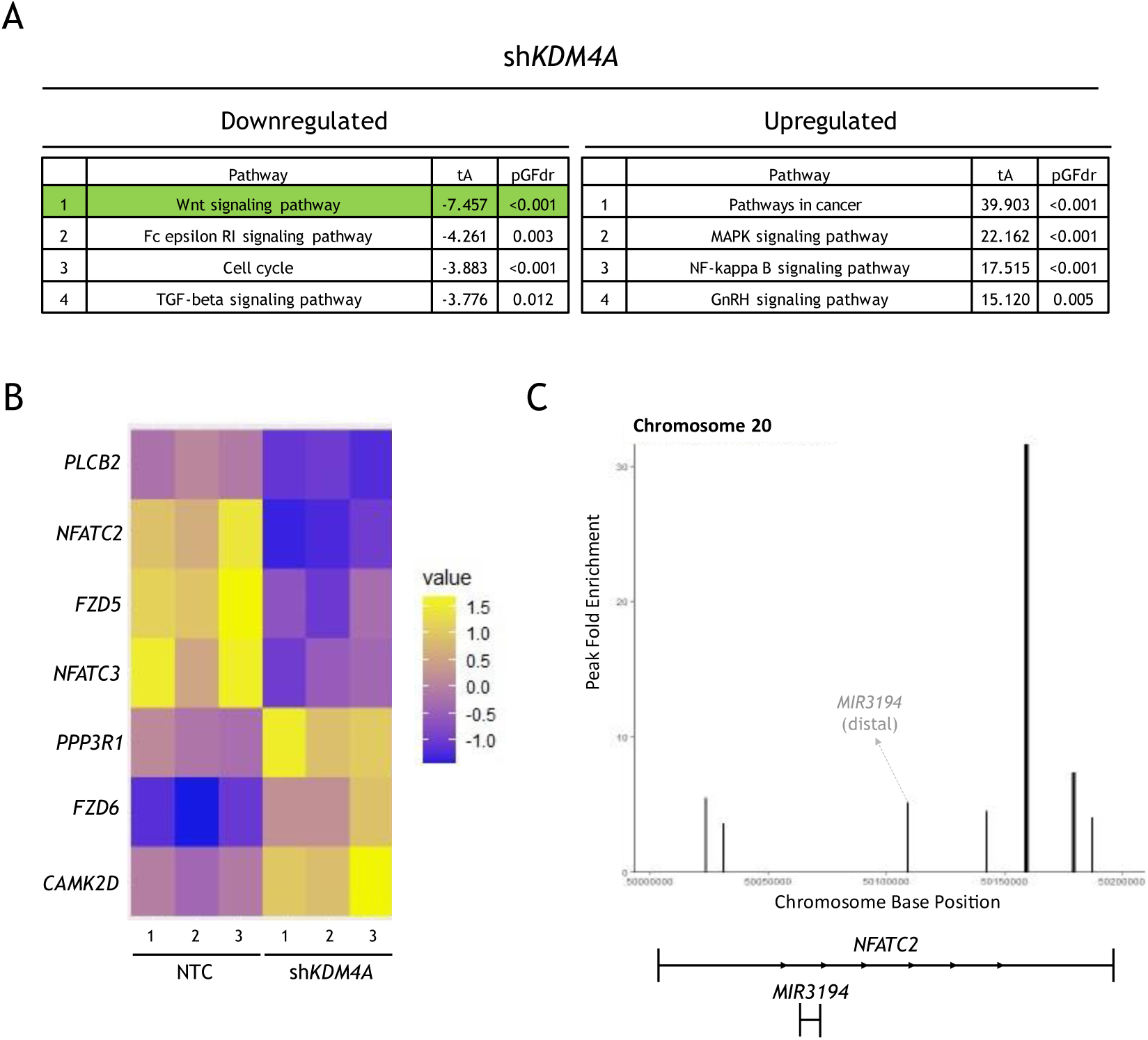
KDM4A transcriptionally regulates *NFATC2* and binds the *NFATC2* gene locus in THP-1 cells. THP-1 cells were transduced with shRNA constructs NTC or sh*KDM4A*. RNA-seq data were generated from cells harvested 48 hr post-puromycin selection (n=3 biological replicates). **(A)** RNA-seq data from transduced THP-1 cells were analysed using the SPIA pathway enrichment tool. Shown are the enrichment statistics (tA and pGFdr) from SPIA for the top five most enriched downregulated and upregulated datasets, for the differential expression between NTC vs. sh*KDM4A*. Pathways with a false discovery rate (FDR/pGFdr) <0.05 were included and ordered by ‘total accumulated perturbation’ (tA). **(B)** The expression of genes from the ‘Wnt signaling pathway’ from SPIA **(A)**, which had p_adj_<0.05 in the RNA-seq dataset for NTC vs sh*KDM4A*, are shown as a heatmap, given as z-scaled FPKM (z-scaling across NTC and two sh*KDM4A* constructs; full heatmap shown in Supplementary Figure 1). **(C)** DNA/protein complexes were immunoprecipitated from untreated THP-1 cells using an anti-KDM4A antibody, and DNA was fragmented and sequenced (n=3 biological replicates). Shown are the KDM4A binding peaks within the *NFATC2* gene region meeting a significance threshold q<0.05, as determined by *epic2*.

Of the possible genes in the SPIA Wnt signaling pathway, seven were significantly deregulated after *KDM4A* KD in THP-1 (Figure 1B) and validated with a second *KDM4A* KD construct (p_adj_<0.05 for both constructs; Supplementary Figure 1). This included downregulation of two members of the NFAT family of transcription factors *NFATC2* and *NFATC3*. Using ChIP-seq peak calls for KDM4A chromatin binding in THP-1 cells (GSE125376), it was also found that KDM4A binds the *NFATC2* gene locus at six sites with statistical significance (Figure 1C; q<0.001). Two of these were within the promoter (<5 kb of the transcription start site (TSS)), indicating that it may regulate *NFATC2* transcription more directly; the remainder were in distal intergenic or intronic regions. One KDM4A binding peak was identified for *NFATC3* (Supplementary Figure 2A), also <5 kb from the TSS, suggesting a potential regulatory role in transcription. Analysis of the TARGET open-source gene expression datasets for paediatric AML patient samples (n=199) showed that *NFATC2* and *NFATC3* expression were correlated (*p*<0.0001; Supplementary Figure 2B). Given the stronger evidence for *NFATC2* as a KDM4A binding target and the correlation with *NFATC3* expression, *NFATC2* was pursued as a transcriptional target of KDM4A of significant interest.

### THP-1 cells are dependent on *NFATC2* expression for survival *in vitro*

Three AML cell lines of diverse cytogenetic origin, THP-1 (*MLL-AF9*, *TP53*^mut^), HL-60 (amplified *MYC*, *TP53*^del^) and MV4-11 (*FLT3*^ITD^, *MLL-AF4*), lost colony-forming capacity after transduction with shRNA targeting *NFATC2* (sh*NFATC2*-1 and sh*NFATC2*-2), compared to an NTC shRNA (Figure 2A; *p*<0.05). In contrast, MOLM-13 (*FLT3*^ITD^, *MLL-AF9*) colony formation was not significantly deregulated by *NFATC2* KD with either construct sh*NFATC2*-1 or sh*NFATC2*-2 (Figure 2A). These data suggest that *NFATC2* is required for AML cell survival in some, but not all, cytogenetic contexts, and independently of *MLL-AF9* expression.

**Figure 2.**
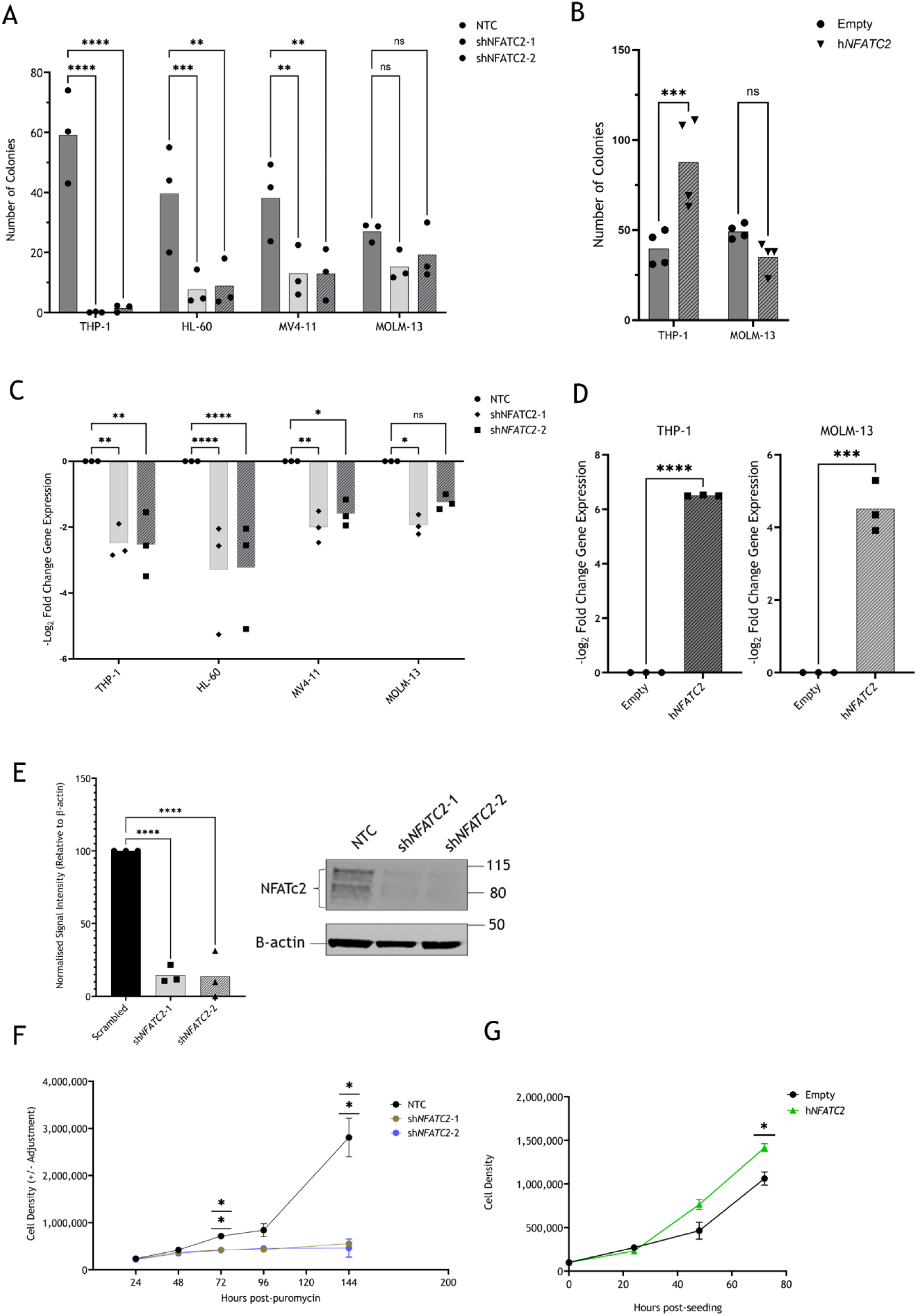
*NFATC2* KD in AML cell lines led to a loss of colony-forming capacity. THP-1 cells were transduced with shRNA constructs NTC, sh*NFATC2*-1 or sh*NFATC2*-2, or either of expression vectors Empty or h*NFATC2*. **(A)** Colony-formation capacity of shRNA-transduced AML cell lines, 7-8 days post-puromycin selection. Mean and individual data replicate numbers of total colonies (n=3 biological replicates). **(B)** Colony-formation capacity of THP-1 or MOLM-13 cells, expressing either Empty or h*NFATC2* vectors, 7-8 days post-seeding. Mean and individual data replicate numbers of total colonies (n=4 biological replicates). **(C)** *NFATC2* expression as measured by qRT-PCR (n=3 biological replicates) in AML cell lines, 48 hr post-puromycin selection. Shown as a mean –log_2_ fold change of expression (with individual replicate points), relative to NTC, using the ΔΔC_t_ method, relative to *BACTIN* and *GAPDH* housekeeping genes. **(D)** *NFATC2* expression as measured by qRT-PCR (n=3 biological replicates) in vector-expressing THP-1 or MOLM-13, presented as in **(C)** and relative to Empty vector. **(E)** Immunoblot of shRNA-transduced THP-1 cells, 72 hr post-puromycin selection. Left: Mean signal intensity (with individual replicate points) as measured by densitometry (n=4 biological replicates). Right: representative immunoblot, with size markers indicated (kDa). **(F)** Mean cell densities (with individual replicate points) of shRNA-transduced THP-1 in liquid culture at intervals post-puromycin selection as shown on the x-axis (n=3 biological replicates). **(G)** Mean cell densities (with individual replicate points) of vector-expressing THP-1 in liquid culture at intervals post-seeding as shown on the x-axis (n=3 biological replicates). Statistical tests used were two-way ANOVA (A-C), two-sided, unpaired t-test (D), one-way ANOVA (E), and repeated measures ANOVA (F-G). p-values shown for ANOVA tests are derived from post-hoc Dunnett’s tests for comparison of treatment with control means. p-values: *<0.05, **<0.01, ***<0.001, ****<0.0001, ns: not significant.

The cell lines in which the effect of *NFATC2* KD on colony formation was the greatest and least—THP-1 and MOLM-13, respectively—were taken forward for further study. Supporting the above data, stable overexpression (OE) of *NFATC2* (h*NFATC2*) increased the colony-forming capacity of THP-1, but not MOLM-13 (Figure 2B, *p*<0.05 for THP-1). Validation of *NFATC2* KD and OE by qRT-PCR is shown in Figures 2C&D, respectively. KD of NFATc2 protein in THP-1 was confirmed by immunoblot (Figure 2E). In parallel, THP-1 expansion in liquid culture was reduced after *NFATC2* KD and increased with h*NFATC2* OE (Figures 2F&G, respectively).

### *NFATC2* KD in THP-1 cells led to *CCND1* downregulation and G1/S transition arrest

Next, we explored the mechanism by which *NFATC2* expression is required for THP-1 cell growth and colony forming potential *in vitro*. Ki-67/DAPI staining of THP-1 cells demonstrated that the proportion of cells in the G1 phase of the cell cycle was increased and the proportion in the G2-M phases was reduced after shRNA *NFATC2* KD, relative to NTC shRNA at 72 hr post-selection. In addition, the proportion of S phase cells was reduced after sh*NFATC2*-1 transduction, indicating a G1/S transition arrest (Figure 3A; *p*<0.05). Reinforcing this, proliferation tracing with CFSE demonstrated a significant reduction in THP-1 proliferation after *NFATC2* KD in the 96-144 hr post-selection time window, with a trend towards reduction between 24 hr-72 hr (Figure 3B; *p*<0.05 for 96-144 hr). Cell surface expression of Annexin V was not significantly increased after *NFATC2* KD at 72 hr or 144 hr post-puromycin selection, suggesting that apoptosis was not a major contributing mechanism for the observed loss of expansion in THP-1 cells after *NFATC2* KD (Supplementary Figure 3).

**Figure 3.**
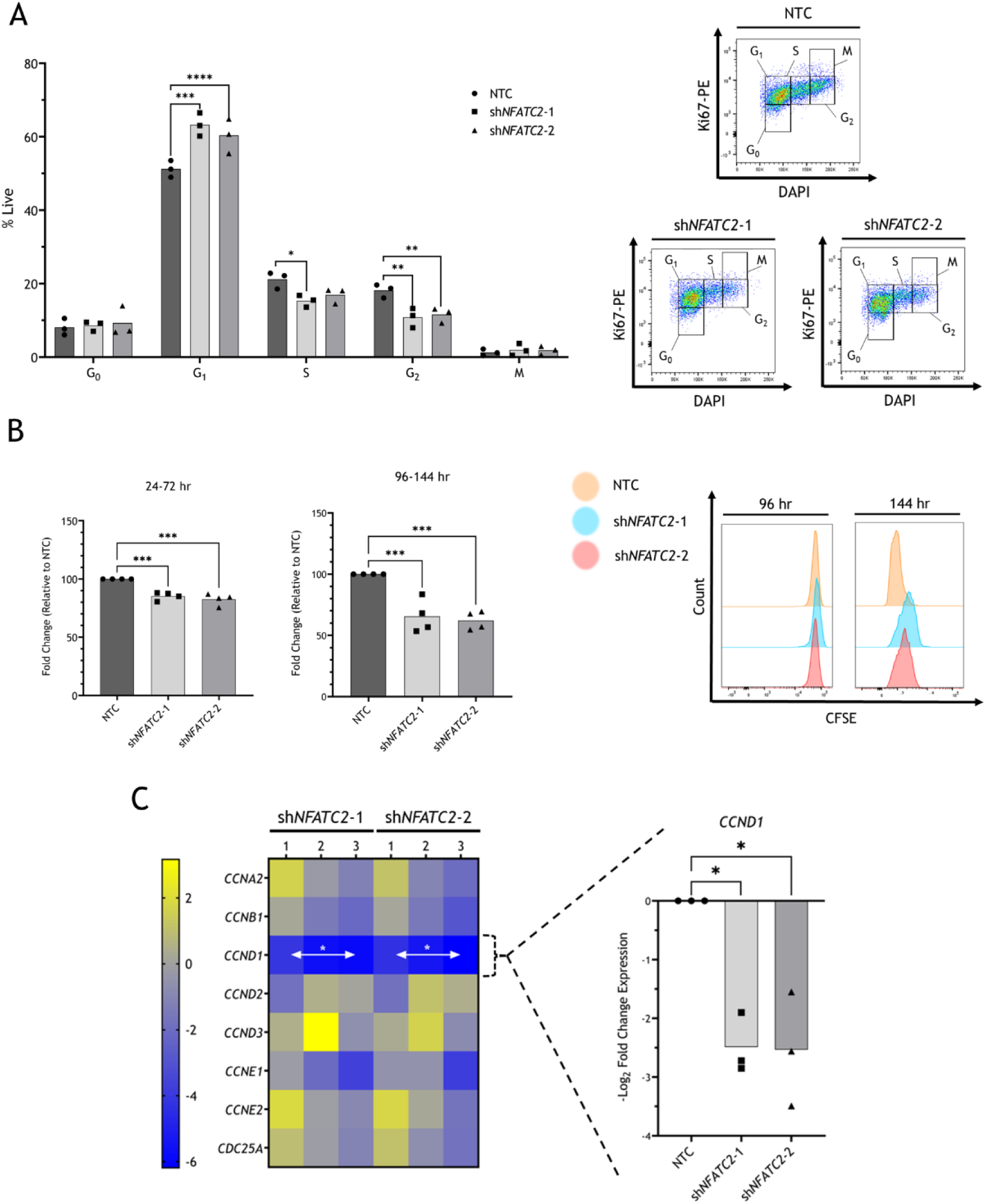
THP-1 cells undergo proliferation arrest after *NFATC2* KD. THP-1 cells were transduced with shRNA constructs NTC, sh*NFATC2*-1 and sh*NFATC2*-2. **(A)** Transduced THP-1 cells stained with Ki67-PE and DAPI and cell cycle phases were quantified by flow cytometry. Left: Mean proportions of cells (with individual replicate points) within each gate after the exclusion of dead cells (n=3 biological replicates). Shown are cells within gates determined to be G_0_, G_1_, S, G_2_ and M phases of the cell cycle. Right: representative flow cytometry plots with gating strategy. **(B)** shRNA-transduced THP-1 cells stained with CFSE for 48 hr (at either 24 or 96 hr post-puromycin selection; n=3 biological replicates) were quantified by flow cytometry. Left: Mean log_2_ fold changes in CFSE intensity of DAPI-negative live cells (with individual replicate points) at the point of staining and 48 hr post-staining. Right: representative flow cytometry histograms showing CFSE signal stained at 96 hr and 48hr later (144 hr timepoint). **(C)** Left: Heatmap showing gene expression data as determined by qRT-PCR from shRNA-transduced THP-1 cells 144 hr post-puromycin selection, for selected genes. Expression is shown as -log_2_ fold changes for each replicate as compared to NTC, using the ΔΔC_t_ method, normalised to *BACTIN* and *GAPDH*. Expression ranges from negative to Right: *CCND1* expression was validated in an independent cohort of shRNA-transduced THP-1 cells (n=3 biological replicates). Shown is *CCND1* expression 144 hr post-puromycin selection as mean -log_2_ fold change (with individual replicate points) compared to NTC using the ΔΔC_t_ method, normalised to *BACTIN* and *GAPDH* housekeeping genes. Statistical tests used were two-way ANOVA (A) and one-way ANOVA (B and C, right), both with post-hoc Dunnett’s tests for comparison of treatment mean against control mean. In C (left), for testing differences of means between NTC (not shown; values are zero) and each treatment group, for each gene, multiple two-sided, unpaired t-tests were used, with correction for multiple testing using the two-stage step-up method of Benjamini, Krieger and Yekutieli. P-values: *<0.05, **<0.01, ***<0.001, ****<0.0001.

Analysis of the expression levels of selected cyclin genes measured at 144 hr post-selection showed that *CCND1* was significantly downregulated after *NFATC2* KD (Figure 3C left; *p*_adj_<0.05). This was validated in a separate cohort of THP-1 cells (Figure 3C right; *p*<0.05), which indicates that cell cycle arrest upon *NFATC2* KD in THP-1 cells is, at least in part, due to *CCND1* downregulation.

### *NFATC2* KD led to global downregulation of *MYC* transcriptional targets in THP-1 cells

RNA-seq was used to reveal the immediate transcriptional targets of *NFATC2* KD, at 24 hr post-selection, relative to NTC shRNA (Figure 4A; GSE173394). GSEA revealed that two of the top pathways deregulated by both shRNA constructs were ‘MYC_TARGETS_V1’ and ‘HALLMARK_OXIDATIVE_PHOSPHORYLATION’ with FDR<0.001 (Figure 4B). The full list of GSEA hallmark pathways which were significantly enriched in the differential genes for NTC vs. sh*NFATC2*-1 and sh*NFATC2*-2 is shown in Supplementary Table 3. These data indicate that *NFATC2* and *MYC* are upstream of some shared gene targets and that *NFATC2* targets are involved in regulating oxidative phosphorylation, amongst other cellular processes.

**Figure 4.**
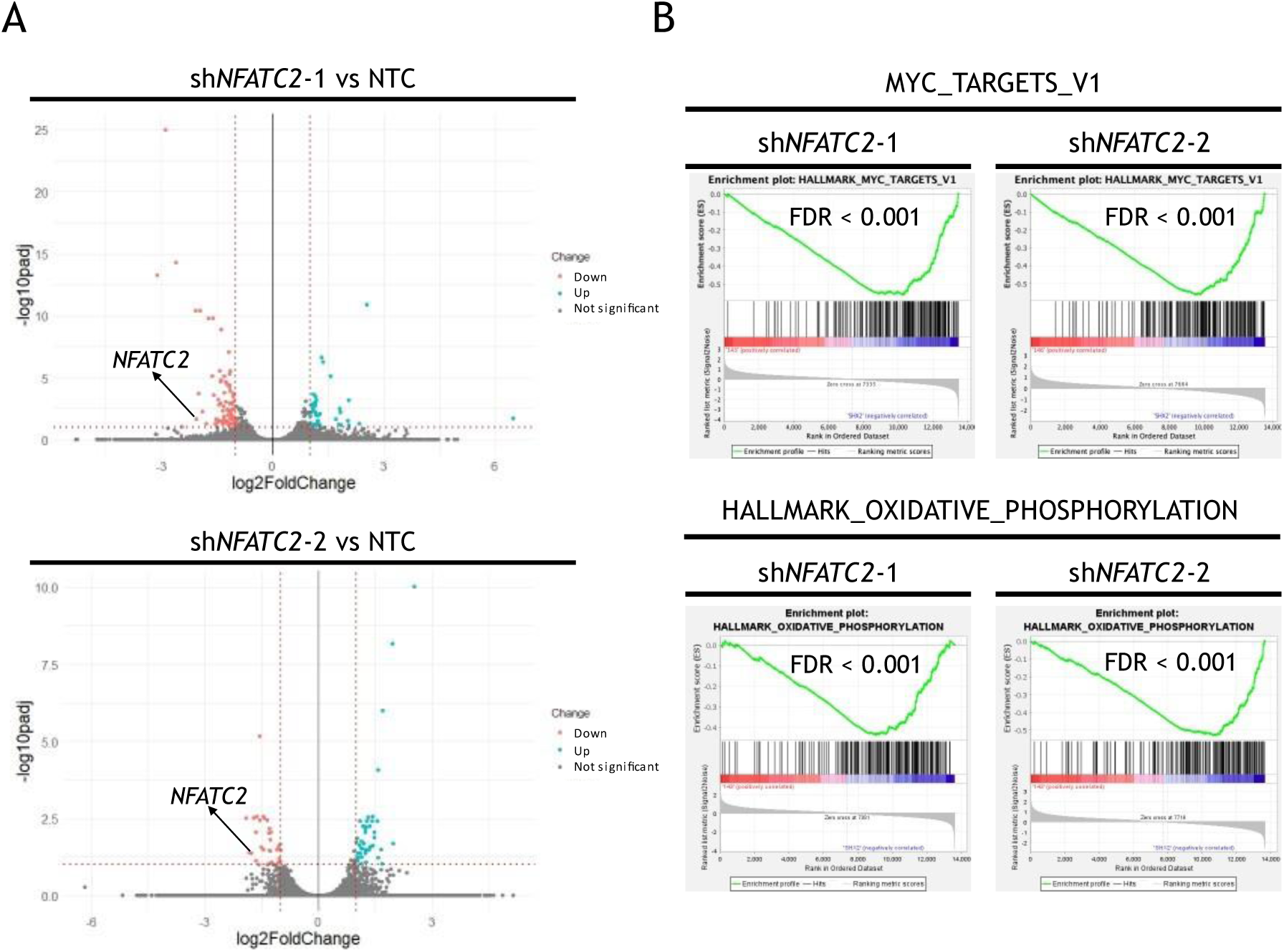
Oxidative phosphorylation and targets of *MYC* are downregulated after *NFATC2* KD in THP-1 cells. THP-1 cells were transduced with shRNA constructs NTC, sh*NFATC2*-1 and sh*NFATC2*-2. RNA-seq data were generated from cells harvested 24 hr post-puromycin selection (n=3 biological replicates). **(A)** Volcano plots show -log_2_ fold changes and -log_10_ p_adj_ values as calculated by DESeq2 for all genes with mean normalised read counts >10. Values are as calculated from 3 pooled replicates. Red dotted lines indicate cut-off thresholds of |-log_2_ fold change| > 1 and -log10 p_adj_ value > 1, with gated genes highlighted in red. **(B)** RNA-seq data from transduced THP-1 cells described in (A) were analysed using the GSEA platform. Shown are the GSEA plots for the datasets HALLMARK_MYC_TARGETS_V1 and HALLMARK_OXIDATIVE_PHOSPHORYLATION, for the differential expression between NTC vs. sh*NFATC2*-1 and sh*NFATC2*-2 as indicated. FDR values are shown.

### NFATc2 protein binds to key mitochondrial genes in THP-1 cells

ChIP-seq analysis for NFATc2 chromatin binding in untreated THP-1 (GSE241260) revealed 517 peaks with statistical significance (FDR<0.05 and log_2_ fold change>1) and 8 peaks were removed due to being in ‘blacklisted’ genome regions. The distribution of NFATc2-bound peaks, relative to the TSS, is shown in Figures 5A&B. GSEA applied to NFATc2-bound peaks showed enrichment of the gene set ‘DANG_BOUND_BY_MYC’ (Figure 5C; FDR=0.05), which is consistent with the finding of ‘MYC_TARGETS_V1’ GSEA enrichment from previous *NFATC2* KD RNA-seq data in THP-1 (Figure 4B). PANTHER analysis of the NFATc2-bound ChIP peaks also revealed that the GO pathway ‘Oxidative phosphorylation’ was enriched (Figure 5D; FDR=0.028), consistent with enrichment of ‘HALLMARK_OXIDATIVE_PHOSPHORYLATION’ from the RNA-seq data analyses. In addition, ChIP peak analysis revealed that the GO pathway ‘Mitochondrial complex IV’ was enriched relative to the input DNA (Figure 5D; DR=0.041), consistent with the enrichment of the GSEA pathway ‘HALLMARK_OXIDATIVE_PHOSPHORYLATION’ from the RNA-seq data analyses.

**Figure 5.**
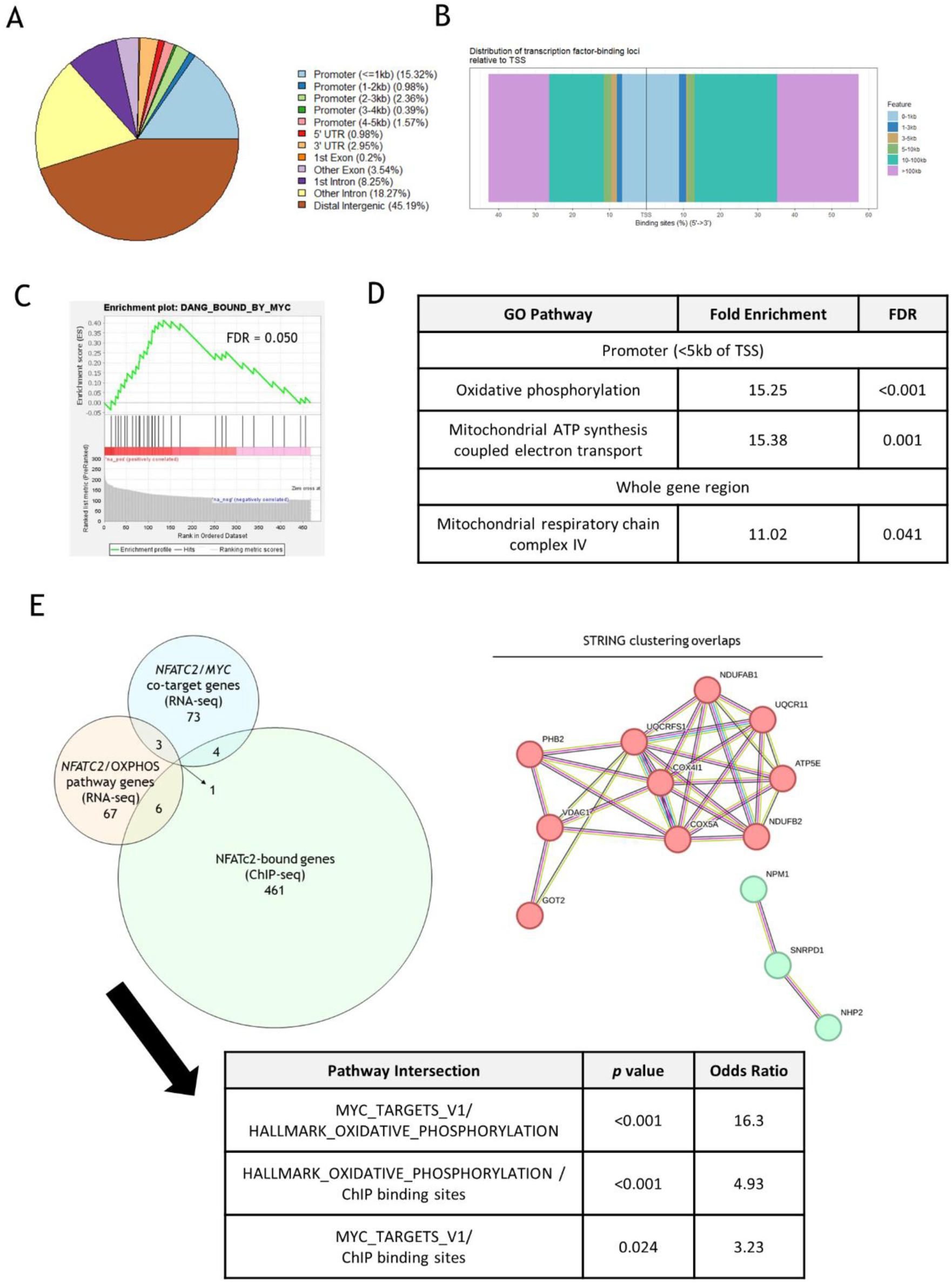
NFATc2 binds genes involved in oxidative phosphorylation and mitochondrial respiration in THP-1 cells. DNA/protein complexes were immunoprecipitated from untreated THP-1 cells using an NFATc2 antibody, and DNA was fragmented and sequenced (n=3 biological replicates). **(A)** Output from the *ChIPseeker* package analysis of NFATc2 binding peaks meeting the significance threshold (FDR<0.05 and log_2_ fold change>1), and without blacklisted regions, as determined by *epic2*. The distribution of binding sites relative to the TSS of the nearest gene (n=138) is given as a pie chart in **(A)**; as a bidirectional plot given in the 5’→3’ direction in **(B)**. **(C)** Protein-coding genes were isolated from the NFATc2-bound *epic2* peaks and analysed using the GSEA platform. Shown is the enrichment plot for the pathway ‘DANG_BOUND_BY_MYC’ with the FDR statistic given. **(D)** Protein-coding genes in (C) were used to find enriched gene ontology (GO) pathways with the PANTHER web tool. Shown are two pathways with the enrichment and FDR statistics. **(E)** Genes from the leading-edge subsets of the pathways enriched in the *NFATC2* RNA-seq KD data, which were present in the subset for both shRNA constructs, and overlapped with NFATc2-bound genes from ChIP-seq analysis; represented by Venn diagram (left). A hypergeometric test was conducted to test pairwise overlaps and the statistics of these are given (bottom). The overlapping genes were also entered into the STRING database and the interaction clusters are also shown (right).

The overlap of genes included in the leading-edge subsets of the enriched GSEA pathways ‘MYC_TARGETS_V1’ and ‘HALLMARK_OXIDATIVE_PHOSPHORYLATION’ after *NFATC2* KD and the ChIP-seq NFATc2-bound genes is shown in Figure 5E (left). These overlaps were statistically significant by hypergeometric test (*p*<0.05; Figure 5E (bottom)), indicating that these genes found from separate analyses are of significance, beyond statistical chance. Of the 13 overlapping genes, the corresponding proteins were found to form 2 clusters using the STRING protein database (Figure 5E, right), of which all 10 proteins in the primary cluster were mitochondrial components. This suggests that key transcriptional targets of *NFATC2*, which are also published targets of *MYC* and components of oxidative phosphorylation, are components of mitochondrial function.

### *MYC* is upstream of *NFATC2* expression

KD of *MYC* in THP-1 cells with either sh*MYC*-1 or sh*MYC*-2 constructs led to a complete loss of colony-forming capacity (Figure 6A; *p*<0.05). *MYC* KD was validated by qRT-PCR (Figure 6B; *p*<0.05). Similar to *NFATC2* KD, Ki-67/DAPI staining of THP-1 cells with *MYC* KD demonstrated that proportions of cells in the G1 phase of the cell cycle were increased and those in the S and G2-M phases were reduced after shRNA *MYC* KD, relative to NTC shRNA at 72 hr post-selection (Figure 6C). Consistent with this observation, proliferation tracing with CTV showed a reduction in the expansion of THP-1 cells after sh*MYC* transduction, relative to NTC (Figure 6D).

**Figure 6.**
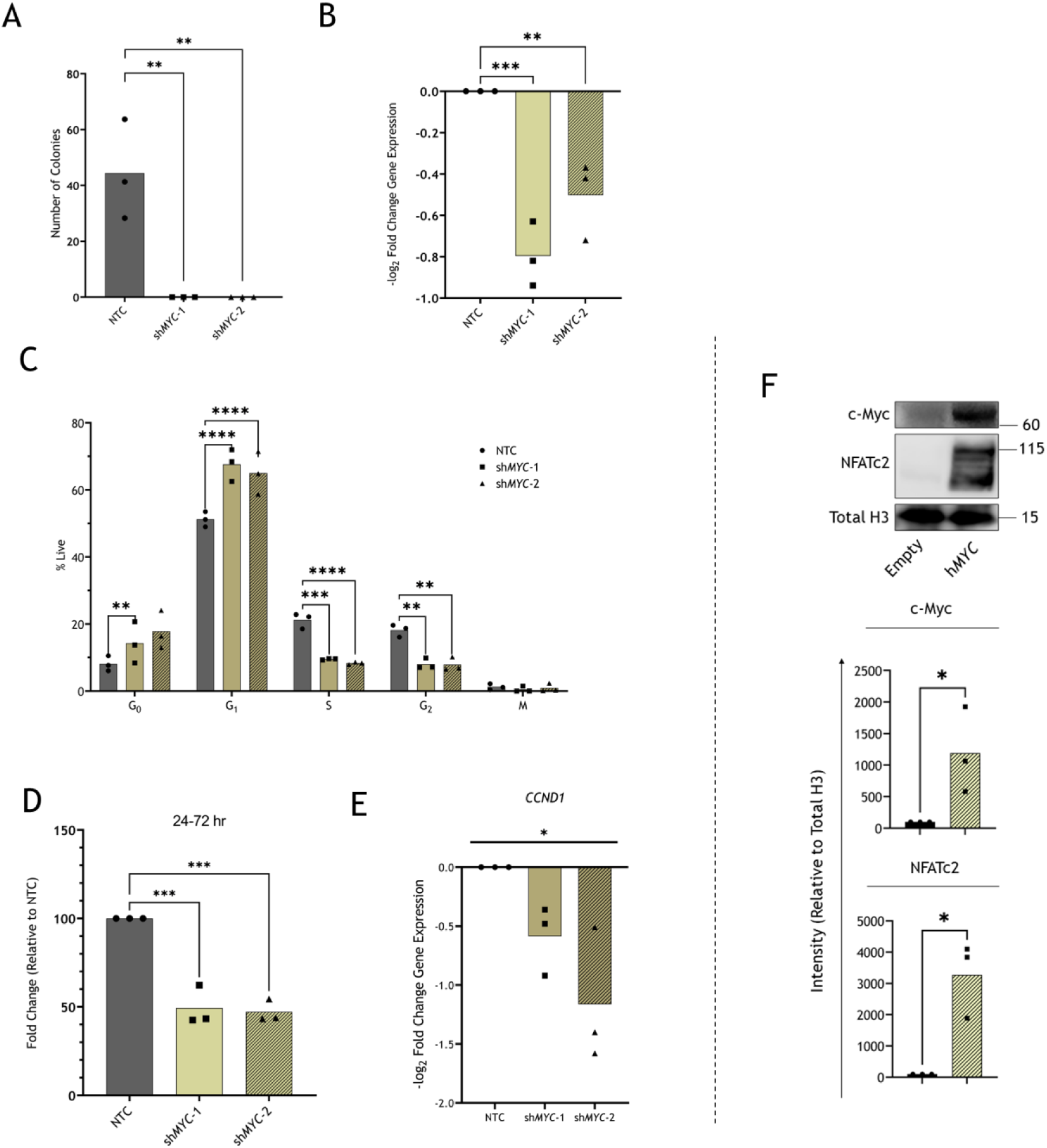
*MYC* KD in THP-1 cells phenocopies KD of *NFATC2*. **(A)** Colony formation capacity of NTC or shMYC-transduced THP-1 cells, 7-8 days post-puromycin selection (n=3 biological replicates). Mean number of total colonies (with individual replicate points). **(B)** *MYC* expression measured in shRNA-transduced THP-1 cells using qRT-PCR at 48 hr post-puromycin selection. Differential expression is shown, as compared between either NTC vs. sh*MYC*-1 and NTC vs. sh*MYC*-2 and are shown as mean -log_2_ fold changes (with individual replicate points; n=3 biological replicates). **(C)** THP-1 cells transduced with shRNA constructs NTC, sh*MYC*-1 and sh*MYC*-2 were stained with Ki67-PE and DAPI (n=3 biological replicates). Mean proportions of cells shown (with individual replicate points) within each gate: G_0_, G_1_, S, G_2_ and M phases of the cell cycle (Figure 3A). **(D)** shRNA-transduced THP-1 cells stained with CTV and 7-AAD with quantification by flow cytometry. Mean log_2_ fold changes in CTV intensity of 7-AAD-negative cells (with individual replicate points) at the point of staining to 48 hr post-staining. Shown are changes from 24 to 72 hr (n=3 biological replicates) post-puromycin selection. **(E)** *CCND1* expression measured in shRNA-transduced THP-1 cells using qRT-PCR at 48 hr post-puromycin selection. Differential expression is shown, as compared between either NTC vs. sh*MYC*-1 and NTC vs. sh*MYC*-2 and are shown as mean -log_2_ fold changes (with individual replicate points; n=3 biological replicates). **(F)** c-Myc and NFATc2 protein expression were measured by immunoblot in THP-1 cells expressing either of the vectors Empty or h*MYC*. Quantitative densitometry results are shown (n=3 biological replicates), expression relative to total histone 3 (H3), and the representative immunoblot is shown with size markers indicated (kDa). The statistic for overall difference between groups is given from a one way ANOVA. Statistical tests used were one-way ANOVA (A, B, D, E), two-way ANOVA (C) and two-sided, unpaired t-tests (A); p-values shown for ANOVA tests are for post-hoc Dunnett’s tests between control and treatment group means (A-D) or the ANOVA test for differences across all groups (E). p-values: *<0.05, **<0.01, ***<0.001, ****<0.0001.

Furthermore, *CCND1* was downregulated after *MYC* KD in THP-1 cells (Figure 6E; *p*<0.05). Together, these data suggest that KD of *MYC* phenocopies that of *NFATC2* in THP-1 cells.

*MYC* (h*MYC*) OE in THP-1 cells was shown to significantly increase the expression of NFATc2 protein (Figure 6F; *p*<0.05) and transcript (Supplementary Figure 4A; *p*<0.05), rendering *MYC* as a plausible transcriptional or translational regulator of NFATc2 expression. Conversely, shRNA *MYC* KD in THP-1 did not lead to significant *NFATC2* downregulation (Supplementary Figure 4B), which could suggest that *NFATC2* expression is not entirely dependent on elevated *MYC* in these cells. In addition, THP-1 cells overexpressing *NFATC2* displayed significantly downregulated c-Myc protein expression, suggesting that there may be a bi-directional relationship (Supplementary Figure 4). Together these data indicate that there is a *MYC–NFATC2* regulatory axis in which *MYC* can upregulate *NFATC2*, but is not necessary for basal *NFATC2* expression, and, in addition, there may be negative feedback from *NFATC2* towards *MYC* in THP-1 cells.

### *NFATC2* maintains oxidative phosphorylation as a mechanism of survival in THP-1 cells

To build on the data from the RNA-seq and ChIP-seq studies, some functional assays were used to elucidate the contribution(s) of *NFATC2* and *MYC* to basic mitochondrial function in THP-1 cells. The specific features of mitochondrial and/or cellular metabolic activity measured by the assays described here are highlighted in Figure 7A.

**Figure 7.**
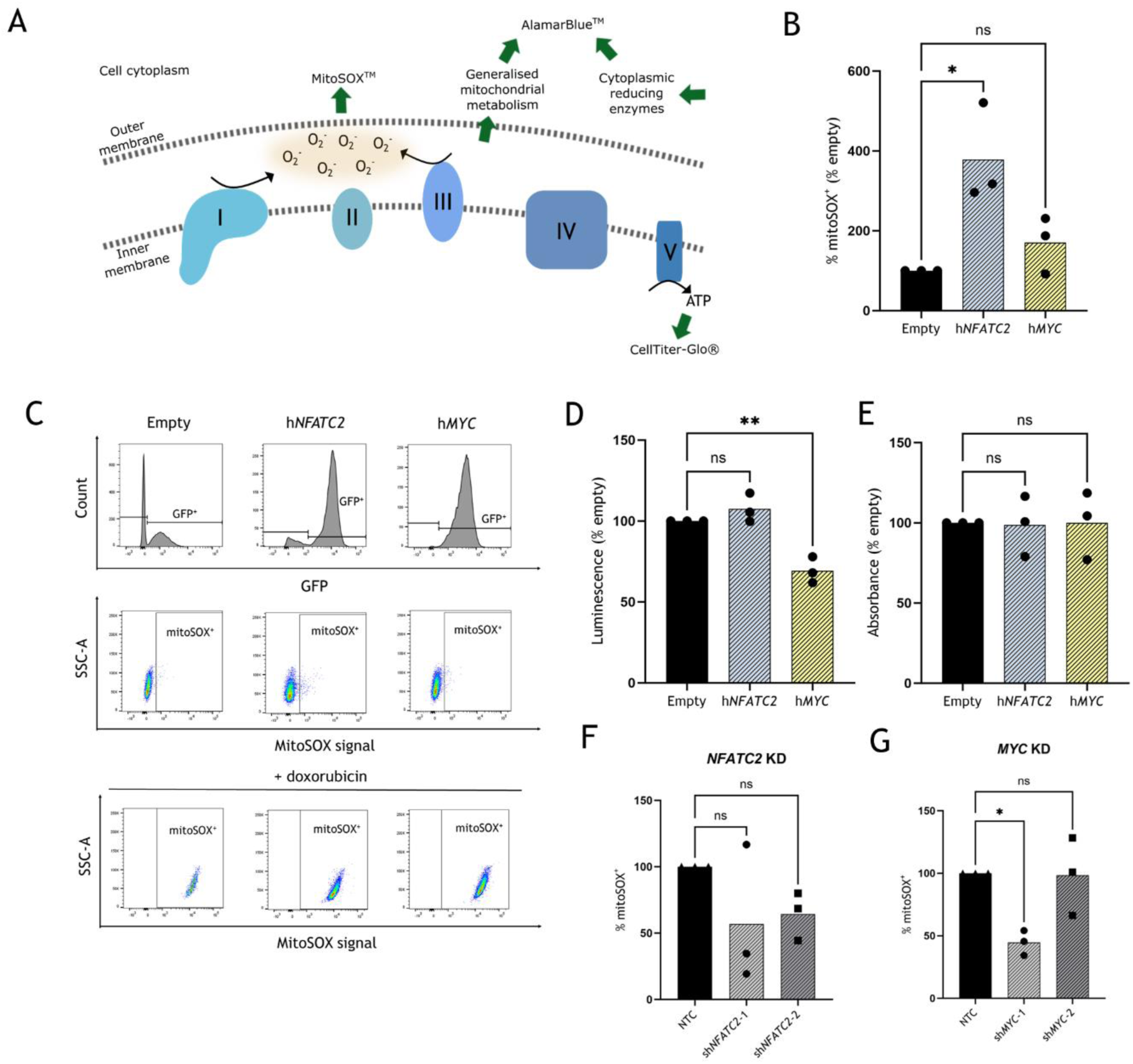
Overexpression of *NFATC2*, but not *MYC*, leads to increased mitochondrial superoxide in THP-1 cells. **(A)** Schematic diagram depicting metabolic activity at the mitochondrial membranes, with complexes I-V indicated. Green arrows highlight points where the distinct assays measure specific outputs of mitochondrial/cellular metabolism, namely: MitoSOX^TM^, measuring mitochondrial superoxide; CellTiter-Glo^®^, measuring ATP; AlamarBlue^TM^, measuring general redox activity from mitochondrial and cellular sources. **(B)** THP-1 cells overexpressing Empty vector, h*NFATC2* or h*MYC* were stained with mitoSOX red dye and DAPI. Viable (DAPI-negative) Mitosox^+^ were quantified initially as a fraction of GFP^+^ cells, and this is shown as mean % Empty vector (with individual replicate points; n=3 biological replicates). **(C)** The flow cytometry gating strategies for MitoSOX assays is shown. Top: gating on the GFP^+^ fraction for each cell line; Middle: gating on mitoSOX^+^ cells; Bottom: gating on MitoSOX^+^ cells using data for cells treated with 5 µM doxorubicin, as a positive control. **(D)** THP-1 cells overexpressing h*NFATC2* or hMYC were incubated with CellTiter-Glo^®^ and luminescence measured after 30 min, which is given as mean % Empty vector (with individual replicate points; n=3 biological replicates). **(E)** THP-1 cells overexpressing h*NFATC2* or h*MYC* were incubated with AlamarBlue^TM^ and absorbance measured after 4 hr, which is given as % Empty vector. **(F-G)** shRNA-transduced THP-1 cells with either NTC, *NFATC2* KD (D) and/or *MYC* KD shRNA were stained with MitoSOX red dye and DAPI. Viable (DAPI-negative) GFP^+^ MitoSOX^+^ cells are quantified as a % GFP^+^ cells and are expressed as mean % Empty vector (with individual replicate points; n=3 biological replicates). p-values: *<0.05, **<0.01, ns: not significant.

MitoSOX^TM^ red fluoresces upon interaction with mitochondrial superoxide, a type of reactive oxygen species (ROS) that arises due to electron leakage from the mitochondrial electron transport chain, and can serve as a marker of mitochondrial respiratory activity. Cells overexpressing h*NFATC2*, but not h*MYC*, had a significantly larger subset of MitoSOX^+^ cells than those expressing the empty vector (Figure 7B). GFP positivity was used to confirm the true expression of the retroviral vector and doxorubicin was applied as a positive control for superoxide release (Figure 7C). These data indicate that h*NFATC2* OE in THP-1 results in a higher basal level of superoxide.

Levels of ATP, which can be a surrogate marker of mitochondrial respiration, were not increased in THP-1 expressing h*NFATC2*, compared to the Empty vector-expressing THP-1, as measured by CellTiter-Glo® in cell suspensions with equivalent densities (Figure 7D). This suggests that the ATP output of mitochondria was not altered by h*NFATC2* OE in THP-1 cells. In contrast, THP-1 expressing h*MYC* had a lower value of CellTiter-Glo®, indicating lower ATP levels (Figure 7D). Using an AlamarBlue^TM^ assay in cell suspensions with equivalent densities, no observable enhancement in resazurin reduction in THP-1 expressing h*NFATC2* or h*MYC* was seen, compared to empty vector (Figure 7E; n=3), indicating similar levels of viability and basal global cellular metabolism.

*NFATC2* KD in THP-1 led to a trend towards reduced MitoSOX^TM^ red staining, but this was below statistical significance (Figure 7F; p_adj_=0.246 and 0.352, for sh*NFATC2*-1 and -2, respectively). *MYC* KD in THP-1 led to reduced MitoSOX^TM^ red staining, but only after transduction with sh*MYC*-1 construct (Figure 7G; p=0.020). Immunoblots for complex IV (COX IV) in THP-1 expression h*NFATC2* or Empty vector, or with NTC/sh*NFATC2* KD, showed no significant changes in quantity, relative to either total H3 or β-actin (Figure 7H and 7I). Since COX IV is a rate-limiting step in mitochondrial electron transport, this finding demonstrates that the changes in superoxide levels with either *NFATC2* OE or KD are not due to altered COX IV protein expression, but some other factor(s).

These data alone do not offer a clear view of the role of *MYC* in this context and further study is required. However, the data do indicate that *NFATC2* regulates processes which contribute to mitochondrial superoxide accumulation, but not overall ATP synthesis or cellular metabolism, nor the expression of rate-limiting COX IV, in THP-1 cells.

## Discussion

In this study, we demonstrated that AML cell lines of diverse cytogenetic backgrounds are dependent on *NFATC2* for colony formation, which, at least in the case of THP-1 cells, is due to *NFATC2* maintenance of the G1/S phase cell cycle transition. The roles of NFATs in cell cycle regulation are well-described; for example, pan-NFAT inhibition using cyclosporine A (CsA) led to increased proliferation of the myeloid compartment [11], indicating a cell cycle-suppressive role for NFATs in these cells. Consistent with this, constitutively active *NFATC2* exhibited a pro-arrest and pro-apoptosis phenotype in NIH-3T3 fibroblasts [17]. This is contrary to our finding, which implies that *NFATC2* maintains cell proliferation in the THP-1 model.

Cyclin D1 is a well-known regulator of the G1-to-S transition in the cell cycle [18, 19] and so our finding of *CCND1* downregulation after *NFATC2* KD is consistent with the arrest phenotype in this context. Elsewhere, NFAT-binding DNA motifs have been characterised within the *CCND1* promoter, and NFATc1, specifically, has been demonstrated to regulate *CCND1-*mediated proliferation in vascular tissues [20-22]; a relationship between *NFATC2* and *CCND1* has not yet been described, to our knowledge. However, *CCND1* downregulation was not significantly demonstrated prior to 144 hr post-puromycin selection, despite *NFATC2* being downregulated at the 48 hr timepoint, which suggests that the *NFATC2–CCND1* relationship is not the result of direct transcriptional activity. To support this, *CCND1* was not found to be an NFATc2-bound gene from the ChIP-seq data and, as such, there are likely to be intermediate regulatory players which we have not characterised here.

The regulation of *CCND1* is complex; its expression and activity are dependent on epigenetic regulation, transcription factors, post-translational modification, and degradation, as covered in several reviews [23-26]. However, in the THP-1 model, we demonstrated that both OE and KD of *NFATC2* either increased or decreased cell expansion, respectively, suggesting that *NFATC2* has a rate-limiting effect on the cell cycle, for which *CCND1* may be a critical factor. In contrast, effects on cell expansion were not observed in MOLM-13 cells with either *NFATC2* OE or KD. While THP-1 and MOLM-13 cells share an MLL-AF9 rearrangement, there are some key mutational differences (e.g., THP-1 carries *FLT3^WT^*, MOLM-13 carries *FLT3^ITD^*), highlighting the importance of the mutational profile-specific context to the observed mechanism(s).

The heterogeneity of NFAT activity is partly due to their interactions with a myriad of transcriptional partners [27], which depends on the cell-specific signalling landscape. For example, transcriptional complexes formed by NFAT(c1-4) and AP-1 heterodimers leads to the integration of calcium and MAPK signalling pathways at the DNA-binding level [28, 29]. Such complexes may also be drug-targetable [30], which could enable the targeting of NFAT complexes in AML cells where a specific combination of signalling pathways is activated. We identified significant enrichment of c-Myc targets within *NFATC2* transcriptional targets, in addition to co-enrichment of c-Myc and NFATc2 DNA-binding targets, via our sequencing experiments. These data suggest that c-Myc could be a transcriptional partner of NFATc2, by which they form DNA-binding complexes to regulate pro-oncogenic signalling, akin to NFAT:AP-1 complexes.

To support this hypothesis, *MYC* KD in THP-1 cells phenocopied *NFATC2* KD, as cells displayed reduced proliferation with evidence of G1/S phase arrest and *CCND1* downregulation, in addition to loss of colony-forming capacity. However, we found that *MYC* OE in THP-1 was accompanied by NFATc2 upregulation, but not vice versa, which suggests that the *NFATC2* is likely a transcriptional target of c-Myc itself; this would be useful to explore with further ChIP-Seq with c-Myc complex precipitation. While our evidence suggests likely DNA co-binding of NFATc2 and c-Myc at common loci, there appears to be a *MYC–NFATC2* regulatory axis which may contribute to the observed effects, and the underlying relationship is more complex.

*MYC* has ubiquitous cellular functions and has been characterised as a pro-leukaemogenic oncogene in AML; for example, it may drive drug resistance [31, 32] and its expression can be prognostic for patient outcome [33]. It can also be targeted therapeutically using inhibition of upstream BET signalling [34]. c-Myc has not been described as a transcriptional partner of NFATc2 previously, although one study has shown NFATc2 to upregulate *MYC* by binding its promoter [35]; while we did not find this from ChIP-seq or *NFATC2* OE experiments, the context of this study was in non-AML cells.

In addition, we identified genes which regulate oxidative phosphorylation (OxPhos) as enriched in *NFATC2-*regulated and NFATc2-bound genes, in THP-1 cells. In support of this, the genes which were in the GSEA leading-edge subsets common to both sh*NFATC2* constructs for the overlaps between ‘MYC_TARGETS_V1’, ‘HALLMARK_OXIDATIVE_PHOSPHORYLATION’, and also ChIP-seq binding sites, were primarily mitochondrial, and clustered together using STRING analysis (Figure 5E).

OxPhos is a highly complex respiratory process with many components (reviewed in [36]) and often cancer cells are described as being more dependent on glycolysis than OxPhos (reviewed in [37]), while some studies have shown AML cells to be vulnerable to inhibition of mitochondria-dependent respiration [38, 39]. We functionally demonstrated that *NFATC2*, but not *MYC*, OE led to an increase of mitochondrial superoxide levels in THP-1, suggesting that there is a role for *NFATC2* in maintaining mitochondrial activity. In addition, it is known that the accumulation of ROS can drive mutagenesis in AML cells (reviewed in [40]), which could support a hypothesis that *NFATC2*-driven superoxide accumulation is an enabling mechanism for further leukaemogenesis and may have a role in AML progression.

The results from *NFATC2* and *MYC* KD are less clear, though there may be a trend towards downregulated superoxide with either KD. From these data, a role for *MYC* in OxPhos cannot be discerned directly, although it may have some input via its maintenance of *NFATC2* expression. Furthermore, despite the enrichment of mitochondrial complex IV genes in the NFATc2 DNA-binding targets, there was no change in complex IV protein quantity observed with either *NFATC2* KD or OE, suggesting that *NFATC2* is not a rate-limiting factor in terms of mitochondrial or, specifically, complex IV synthesis.

Clearly the role(s) of *NFATC2* and *MYC* in OxPhos and, more generally, mitochondrial function require further characterisation in future study, and these data underpin a rationale for doing so. THP-1 cells have been demonstrated to be highly OxPhos-dependent [41], and so this model may be biased; as such, further study should expand beyond this model alone. In addition, KDM4A enzymatic activity is known to be highly oxygen sensitive [42], which could imply that the *KDM4A*–*NFATC2* axis has specific roles (or a lack thereof) to play across the variably hypoxic environments within the bone marrow microenvironment, at the site of AML residual disease; again, more sophisticated *in vitro* models should be used to assess this further.

Inhibition of NFAT activation is well-established pre-clinically and clinically using drugs such as CsA, tacrolimus and VIVIT peptide, but these drugs have been associated with significant toxicity, in part due to their non-specificity towards individual NFAT isoforms [43-45]. There are experimental compounds which can target individual NFAT isoforms by binding specific domains (Kitamura and Kaminuma, 2021) [46], but these are in their infancy. As such, multi-pathway inhibition at the level of the NFAT transcriptional complex—such as the putative NFATc2–c-Myc complex—or dual upstream pathway inhibition may enable the administration of sub-toxic doses, while retaining specificity to cells with such an active oncogenic mechanism.

Consider also that this study was conducted primarily in THP-1, a cell line derived from infant AML with MLL-AF9, meaning that the results are not generalisable to patients from older cohorts or with differing molecular profiles. Although *NFATC2* KD impaired colony formation in two other models (HL-60 and MV4-11), the specific mechanism was not investigated here. In addition, the effects of *NFATC2* depletion in healthy haemopoietic stem cells should be investigated further, to determine its role in these cells specifically and the potential for a therapeutic window.

Finally, we should consider the role of KDM4A in the mechanisms identified; whether it, too, regulates OxPhos/mitochondrial function and/or *CCND1*, so as to identify the comparative specificity of its inhibition against that of the NFAT family. While a specific inhibitor for NFATc2 is not currently available, there are more specific KDM4 family inhibitors. Alternatively, there may be specific epigenetic markings which correspond to *NFATC2* transcription in AML specifically, and these should be profiled more extensively, to further identify leukaemogenic epigenetic rewiring which may represent points of therapeutic vulnerability.

## Conclusion

In this study, we have identified *NFATC2* as a novel player in AML cell pathobiology, downstream of the histone demethylase *KDM4A,* which has previously been characterised as a master regulator of oncogenesis in AML cells, and highlighted the role of *MYC* as a potential co-regulatory gene. In addition, we have started delineating some putative mechanisms of *NFATC2*-led oncogenesis in this context, which opens avenues for further mechanistic studies and the development of novel therapeutic strategies for AML. Firstly, these cells were found to be dependent on *NFATC2* expression for proliferation, which was mediated by G1/S cell cycle transition and, at least in part, by *CCND1* expression. Secondly, *NFATC2* and *MYC* shared significantly enriched sets of transcriptional and DNA-binding targets, and NFATc2 protein was found to be downstream of *MYC* expression. Thirdly, oxidative phosphorylation genes were enriched in *NFATC2* transcriptional and NFATc2-binding targets, and a subset of mitochondrial genes were common to *MYC* targets also. Of note, *MYC* KD phenocopied *NFATC2* KD in THP-1 cells, together suggesting shared genomic and functional properties. However, while *NFATC2* expression was found to be linked with mitochondrial superoxide levels in these cells, this was not found for *MYC* expression, indicating divergent roles in the nuanced regulation of metabolic activity. Together, these data provide a foundation for further investigation of the novel role(s) of *NFATC2*, its relationship with *MYC*, and dual-targeting possibilities, in the context of other models of AML.

## Cell line authentication

To authenticate cell lines, these were cultured at the recommended conditions and growth curves monitored by cell counts, with a comparison to the supplier’s suggested doubling time. Morphology was also examined under an inverted-light microscope and, after methanol fixation on glass slides, with May–Grünwald–Giemsa staining, and compared with supplier-suggested morphology.

## Data accessibility

The data that support the findings of this study are available on request from the corresponding authors. Sequencing data have been deposited in the GEO repository (NCBI) under the following datasets: GSE125375, ChIP-seq for KDM4A binding in THP-1 cells; GSE125376 RNA-seq for *KDM4A* KD in THP-1 cells; GSE173394, RNA-seq for *NFATC2* KD in THP-1 cells; GSE241260, ChIP-seq for NFATc2 binding in THP-1 cells. The TARGET-AML dataset referenced in Supplementary Figure 2 is available from https://portal.gdc.cancer.gov/projects/TARGET-AML.

## Acknowledgements

We would like to thank the Wellcome Trust [105614/Z/14/Z], the MRC CiC [18048], the Howat Foundation and Friends of the Paul O’Gorman Leukaemia Research Centre, and the Carnegie Trust [PHD007721] for funding this project. In addition, we offer thanks to our colleagues at the Paul O’Gorman Leukaemia Research Centre for technical support, including Jennifer Cassels, who supported FACS work.

## Author contributions

SDP, MEM and XH designed the study; SDP and MEM conducted the experimental work and analysed the data; all authors wrote and edited the manuscript; all authors approved the final manuscript.

## Disclosure/conflict of interest

There are no conflicts of interest to declare.

## Notes

https://www.ncbi.nlm.nih.gov/geo/query/acc.cgi?acc=GSE125375

https://www.ncbi.nlm.nih.gov/geo/query/acc.cgi?acc=GSE125376

https://www.ncbi.nlm.nih.gov/geo/query/acc.cgi?acc=GSE173394

https://portal.gdc.cancer.gov/projects/TARGET-AML

### Competing Interest Statement

The authors have declared no competing interest.

## References

1. (HMRN) HMRN, 2022. Survival statistics: Acute myeloid leukaemia. Haematological Malignancy Research Network (HMRN).

2. Sasaki K, Ravandi F, Kadia TM, DiNardo CD, Short NJ, Borthakur G, Jabbour E & Kantarjian HM (2021) De novo acute myeloid leukemia: A population-based study of outcome in the United States based on the Surveillance, Epidemiology, and End Results (SEER) database, 1980 to 2017. Cancer 127, 2049-2061, doi: 10.1002/cncr.33458.

3. Kantarjian H, Kadia T, DiNardo C, Daver N, Borthakur G, Jabbour E, Garcia-Manero G, Konopleva M & Ravandi F (2021) Acute myeloid leukemia: current progress and future directions. Blood Cancer Journal 11, 41, doi: 10.1038/s41408-021-00425-3.

4. Corces-Zimmerman MR, Hong WJ, Weissman IL, Medeiros BC & Majeti R (2014) Preleukemic mutations in human acute myeloid leukemia affect epigenetic regulators and persist in remission. Proc Natl Acad Sci U S A 111, 2548–2553, doi: 10.1073/pnas.1324297111.

5. Massett ME, Monaghan L, Patterson S, Mannion N, Bunschoten RP, Hoose A, Marmiroli S, Liskamp RMJ, Jørgensen HG, Vetrie D, Michie AM & Huang X (2021) A KDM4A-PAF1-mediated epigenomic network is essential for acute myeloid leukemia cell self-renewal and survival. Cell Death Dis 12, 573, doi: 10.1038/s41419-021-03738-0.

6. Boila LD, Chatterjee SS, Banerjee D & Sengupta A (2018) KDM6 and KDM4 histone lysine demethylases emerge as molecular therapeutic targets in human acute myeloid leukemia. Exp Hematol 58, 44–51.e47, doi: 10.1016/j.exphem.2017.10.002.

7. Harris WJ, Huang X, Lynch JT, Spencer GJ, Hitchin JR, Li Y, Ciceri F, Blaser JG, Greystoke BF, Jordan AM, Miller CJ, Ogilvie DJ & Somervaille TC (2012) The histone demethylase KDM1A sustains the oncogenic potential of MLL-AF9 leukemia stem cells. Cancer Cell 21, 473–487, doi: 10.1016/j.ccr.2012.03.014.

8. Chow CW, Rincón M & Davis RJ (1999) Requirement for transcription factor NFAT in interleukin-2 expression. Mol Cell Biol 19, 2300–2307, doi: 10.1128/mcb.19.3.2300.

9. Kiani A, Habermann I, Haase M, Feldmann S, Boxberger S, Sanchez-Fernandez MA, Thiede C, Bornhäuser M & Ehninger G (2004) Expression and regulation of NFAT (nuclear factors of activated T cells) in human CD34+ cells: down-regulation upon myeloid differentiation. J Leukoc Biol 76, 1057–1065, doi: 10.1189/jlb.0404259.

10. Kiani A, Kuithan H, Kuithan F, Kyttälä S, Habermann I, Temme A, Bornhäuser M & Ehninger G (2007) Expression analysis of nuclear factor of activated T cells (NFAT) during myeloid differentiation of CD34+ cells: regulation of Fas ligand gene expression in megakaryocytes. Exp Hematol 35, 757–770, doi: 10.1016/j.exphem.2007.02.001.

11. Fric J, Lim CX, Koh EG, Hofmann B, Chen J, Tay HS, Mohammad Isa SA, Mortellaro A, Ruedl C & Ricciardi-Castagnoli P (2012) Calcineurin/NFAT signalling inhibits myeloid haematopoiesis. EMBO Mol Med 4, 269–282, doi: 10.1002/emmm.201100207.

12. Fric J, Lim CX, Mertes A, Lee BT, Viganò E, Chen J, Zolezzi F, Poidinger M, Larbi A, Strobl H, Zelante T & Ricciardi-Castagnoli P (2014) Calcium and calcineurin-NFAT signaling regulate granulocyte-monocyte progenitor cell cycle via Flt3-L. Stem Cells 32, 3232–3244, doi: 10.1002/stem.1813.

13. Gregory MA, Phang TL, Neviani P, Alvarez-Calderon F, Eide CA, O’Hare T, Zaberezhnyy V, Williams RT, Druker BJ, Perrotti D & Degregori J (2010) Wnt/Ca2+/NFAT signaling maintains survival of Ph+ leukemia cells upon inhibition of Bcr-Abl. Cancer Cell 18, 74–87, doi: 10.1016/j.ccr.2010.04.025.

14. Metzelder SK, Michel C, von Bonin M, Rehberger M, Hessmann E, Inselmann S, Solovey M, Wang Y, Sohlbach K, Brendel C, Stiewe T, Charles J, Ten Haaf A, Ellenrieder V, Neubauer A, Gattenlöhner S, Bornhäuser M & Burchert A (2015) NFATc1 as a therapeutic target in FLT3-ITD-positive AML. Leukemia 29, 1470–1477, doi: 10.1038/leu.2015.95.

15. Solovey M, Wang Y, Michel C, Metzeler KH, Herold T, Göthert JR, Ellenrieder V, Hessmann E, Gattenlöhner S, Neubauer A, Pavlinic D, Benes V, Rupp O & Burchert A (2019) Nuclear factor of activated T-cells, NFATC1, governs FLT3(ITD)-driven hematopoietic stem cell transformation and a poor prognosis in AML. J Hematol Oncol 12, 72, doi: 10.1186/s13045-019-0765-y.

16. Pear WS, Nolan GP, Scott ML & Baltimore D (1993) Production of high-titer helper-free retroviruses by transient transfection. Proc Natl Acad Sci U S A 90, 8392–8396, doi: 10.1073/pnas.90.18.8392.

17. Robbs BK, Cruz AL, Werneck MB, Mognol GP & Viola JP (2008) Dual roles for NFAT transcription factor genes as oncogenes and tumor suppressors. Mol Cell Biol 28, 7168–7181, doi: 10.1128/mcb.00256-08.

18. Baldin V, Lukas J, Marcote MJ, Pagano M & Draetta G (1993) Cyclin D1 is a nuclear protein required for cell cycle progression in G1. Genes Dev 7, 812–821, doi: 10.1101/gad.7.5.812.

19. Hitomi M & Stacey DW (1999) Cyclin D1 production in cycling cells depends on ras in a cell-cycle-specific manner. Curr Biol 9, 1075–1084, doi: 10.1016/s0960-9822(99)80476-x.

20. Guo ZY, Hao XH, Tan FF, Pei X, Shang LM, Jiang XL & Yang F (2011) The elements of human cyclin D1 promoter and regulation involved. Clin Epigenetics 2, 63–76, doi: 10.1007/s13148-010-0018-y.

21. Karpurapu M, Wang D, Van Quyen D, Kim TK, Kundumani-Sridharan V, Pulusani S & Rao GN (2010) Cyclin D1 is a bona fide target gene of NFATc1 and is sufficient in the mediation of injury-induced vascular wall remodeling. J Biol Chem 285, 3510–3523, doi: 10.1074/jbc.M109.063727.

22. Kundumani-Sridharan V, Singh NK, Kumar S, Gadepalli R & Rao GN (2013) Nuclear Factor of Activated T Cells c1 Mediates p21-activated Kinase 1 Activation in the Modulation of Chemokine-induced Human Aortic Smooth Muscle Cell F-actin Stress Fiber Formation, Migration, and Proliferation and Injury-induced Vascular Wall Remodeling*. Journal of Biological Chemistry 288, 22150–22162, doi: 10.1074/jbc.M113.454082.

23. Alao JP (2007) The regulation of cyclin D1 degradation: roles in cancer development and the potential for therapeutic invention. Mol Cancer 6, 24, doi: 10.1186/1476-4598-6-24.

24. Klein EA & Assoian RK (2008) Transcriptional regulation of the cyclin D1 gene at a glance. J Cell Sci 121, 3853–3857, doi: 10.1242/jcs.039131.

25. Montalto FI & De Amicis F (2020) Cyclin D1 in Cancer: A Molecular Connection for Cell Cycle Control, Adhesion and Invasion in Tumor and Stroma. Cells 9, doi: 10.3390/cells9122648.

26. Pawlonka J, Rak B & Ambroziak U (2021) The regulation of cyclin D promoters – review. Cancer Treatment and Research Communications 27, 100338, doi: 10.1016/j.ctarc.2021.100338.

27. Gabriel CH, Gross F, Karl M, Stephanowitz H, Hennig AF, Weber M, Gryzik S, Bachmann I, Hecklau K, Wienands J, Schuchhardt J, Herzel H, Radbruch A, Krause E & Baumgrass R (2016) Identification of Novel Nuclear Factor of Activated T Cell (NFAT)-associated Proteins in T Cells. J Biol Chem 291, 24172–24187, doi: 10.1074/jbc.M116.739326.

28. Chen L, Glover JN, Hogan PG, Rao A & Harrison SC (1998) Structure of the DNA-binding domains from NFAT, Fos and Jun bound specifically to DNA. Nature 392, 42–48, doi: 10.1038/32100.

29. Lopez-Rodríguez C, Aramburu J, Rakeman AS & Rao A (1999) NFAT5, a constitutively nuclear NFAT protein that does not cooperate with Fos and Jun. Proc Natl Acad Sci U S A 96, 7214–7219, doi: 10.1073/pnas.96.13.7214.

30. Mognol GP, González-Avalos E, Ghosh S, Spreafico R, Gudlur A, Rao A, Damoiseaux R & Hogan PG (2019) Targeting the NFAT:AP-1 transcriptional complex on DNA with a small-molecule inhibitor. Proc Natl Acad Sci U S A 116, 9959–9968, doi: 10.1073/pnas.1820604116.

31. Fauriat C & Olive D (2014) AML drug resistance: c-Myc comes into play. Blood 123, 3528–3530, doi: 10.1182/blood-2014-04-566711.

32. Li L, Osdal T, Ho Y, Chun S, McDonald T, Agarwal P, Lin A, Chu S, Qi J, Li L, Hsieh YT, Dos Santos C, Yuan H, Ha TQ, Popa M, Hovland R, Bruserud Ø, Gjertsen BT, Kuo YH, Chen W, Lain S, McCormack E & Bhatia R (2014) SIRT1 activation by a c-MYC oncogenic network promotes the maintenance and drug resistance of human FLT3-ITD acute myeloid leukemia stem cells. Cell Stem Cell 15, 431–446, doi: 10.1016/j.stem.2014.08.001.

33. Ohanian M, Rozovski U, Kanagal-Shamanna R, Abruzzo LV, Loghavi S, Kadia T, Futreal A, Bhalla K, Zuo Z, Huh YO, Post SM, Ruvolo P, Garcia-Manero G, Andreeff M, Kornblau S, Borthakur G, Hu P, Medeiros LJ, Takahashi K, Hornbaker MJ, Zhang J, Nogueras-González GM, Huang X, Verstovsek S, Estrov Z, Pierce S, Ravandi F, Kantarjian HM, Bueso-Ramos CE & Cortes JE (2019) MYC protein expression is an important prognostic factor in acute myeloid leukemia. Leuk Lymphoma 60, 37–48, doi: 10.1080/10428194.2018.1464158.

34. Abraham SA, Hopcroft LEM, Carrick E, Drotar ME, Dunn K, Williamson AJK, Korfi K, Baquero P, Park LE, Scott MT, Pellicano F, Pierce A, Copland M, Nourse C, Grimmond SM, Vetrie D, Whetton AD & Holyoake TL (2016) Dual targeting of p53 and c-MYC selectively eliminates leukaemic stem cells. Nature 534, 341–346, doi: 10.1038/nature18288.

35. Mognol GP, de Araujo-Souza PS, Robbs BK, Teixeira LK & Viola JP (2012) Transcriptional regulation of the c-Myc promoter by NFAT1 involves negative and positive NFAT-responsive elements. Cell Cycle 11, 1014–1028, doi: 10.4161/cc.11.5.19518.

36. Smeitink J, van den Heuvel L & DiMauro S (2001) The genetics and pathology of oxidative phosphorylation. Nature Reviews Genetics 2, 342–352, doi: 10.1038/35072063.

37. Zheng J (2012) Energy metabolism of cancer: Glycolysis versus oxidative phosphorylation (Review). Oncol Lett 4, 1151–1157, doi: 10.3892/ol.2012.928.

38. Skrtić M, Sriskanthadevan S, Jhas B, Gebbia M, Wang Z, Hurren R, Jitkova Y, Gronda M, Maclean N, Lai CK, Eberhard Y, Bartoszko J, Spagnuolo P, Rutledge AC, Datti A, Ketela T, Moffat J, Robinson BH, Cameron JH, Wrana J, Eaves CJ, Minden MD, Wang JC, Dick JE, Humphries K, Nislow C, Giaever G & Schimmer AD (2011) Inhibition of mitochondrial translation as a therapeutic strategy for human acute myeloid leukemia. Cancer Cell 20, 674–688, doi: 10.1016/j.ccr.2011.10.015.

39. Sriskanthadevan S, Jeyaraju DV, Chung TE, Prabha S, Xu W, Skrtic M, Jhas B, Hurren R, Gronda M, Wang X, Jitkova Y, Sukhai MA, Lin FH, Maclean N, Laister R, Goard CA, Mullen PJ, Xie S, Penn LZ, Rogers IM, Dick JE, Minden MD & Schimmer AD (2015) AML cells have low spare reserve capacity in their respiratory chain that renders them susceptible to oxidative metabolic stress. Blood 125, 2120–2130, doi: 10.1182/blood-2014-08-594408.

40. Sallmyr A, Fan J & Rassool FV (2008) Genomic instability in myeloid malignancies: Increased reactive oxygen species (ROS), DNA double strand breaks (DSBs) and error-prone repair. Cancer Letters 270, 1–9, doi: 10.1016/j.canlet.2008.03.036.

41. Suganuma K, Miwa H, Imai N, Shikami M, Gotou M, Goto M, Mizuno S, Takahashi M, Yamamoto H, Hiramatsu A, Wakabayashi M, Watarai M, Hanamura I, Imamura A, Mihara H & Nitta M (2010) Energy metabolism of leukemia cells: glycolysis versus oxidative phosphorylation. Leukemia & Lymphoma 51, 2112–2119, doi: 10.3109/10428194.2010.512966.

42. Hancock RL, Masson N, Dunne K, Flashman E & Kawamura A (2017) The Activity of JmjC Histone Lysine Demethylase KDM4A is Highly Sensitive to Oxygen Concentrations. ACS Chemical Biology 12, 1011–1019, doi: 10.1021/acschembio.6b00958.

43. Klintmalm G, Sundelin B, Bohman S-O & Wilczek H (1984) INTERSTITIAL FIBROSIS IN RENAL ALLOGRAFTS AFTER 12 TO 46 MONTHS OF CYCLOSPORIN TREATMENT: BENEFICIAL EFFECT OF LOW DOSES IN EARLY POST-TRANSPLANTATION PERIOD. The Lancet 324, 950–954, doi: 10.1016/S0140-6736(84)91166-8.

44. Noguchi H, Matsushita M, Okitsu T, Moriwaki A, Tomizawa K, Kang S, Li ST, Kobayashi N, Matsumoto S, Tanaka K, Tanaka N & Matsui H (2004) A new cell-permeable peptide allows successful allogeneic islet transplantation in mice. Nat Med 10, 305–309, doi: 10.1038/nm994.

45. Tedesco D & Haragsim L (2012) Cyclosporine: a review. J Transplant 2012, 230386, doi: 10.1155/2012/230386.

46. Kitamura N & Kaminuma O (2021) Isoform-Selective NFAT Inhibitor: Potential Usefulness and Development. Int J Mol Sci 22, doi: 10.3390/ijms22052725.

