## Supplementary Files for "*MYC–NFATC2 axis* maintains cell cycle and mitochondrial function in AML cells"

### Supplementary Methods

#### *Plasmid preparation*

Stbl3™ chemically competent *E. coli* were transformed with lentiviral and retroviral plasmid vectors (Addgene, Sigma-Aldrich) and subsequently expanded using LB agar plates and Terrific broth, with ampicillin (100 µg/mL; Merck) selection. Plasmids were extracted using a QIAprep Spin Miniprep Kit or Plasmid Maxi Kit (Qiagen) as per the manufacturer's instructions. Expected plasmid sequences were confirmed using restriction digestion with 0.8% (W/V) agarose gel electrophoresis and DNA sequencing (Eurofins Genomics).

#### *Flow cytometry*

All flow cytometry assays were conducted on a FACSCanto™ II flow cytometer and data were acquired using BD FACSDiva™ software, with analyses conducted on FlowJo software (v10.7.2). Apoptosis: Cells were incubated with Annexin V conjugated to FITC or APC (1:100 dilution in HBSS; BD Biosciences) and DAPI (1 µg/mL; Thermo Fisher Scientific) for 15 min at RT. Using this staining, cells were gated into either Annexin$^+$ or Annexin$^-$ fractions.

Cells were stained with CFSE (CarboxyFluoroscein Succinimidyl Ester) and CTV (CellTrace™ Violet; both Thermo Fisher Scientific) at 5 µM in the appropriate base medium for the desired cell line and incubated at 37°C with 5% CO2 for 20 min, before staining with DAPI (1 µg/mL), washing and subsequent data acquisition. The geometric mean for CFSE (in the 488-530/30 channel) was established at the point of staining and subsequent timepoints.

In the Ki67/DAPI assay, harvested cells were washed and fixed in cold 70% ethanol before overnight storage at -20°C. Cells were then washed in PBS/1% (V/V) FBS and stained with anti-Ki-67-PE (1:100 dilution; BD) for 30 min at RT, prior to 5 min incubation with DAPI (1 µg/mL;). Using the Ki-67-PE and DAPI flow cytometry distribution, cells were categorised into the following phases of the cell cycle: G0/G1, S, G2, and M.

*RNA extraction, cDNA synthesis, and qRT-PCR*

RNA was extracted using an RNeasy® Mini Plus Kit or an RNeasy® Micro Plus Kit (Qiagen). cDNA was synthesised from RNA using a SuperScript™ IV Reverse Transcriptase first-strand synthesis kit (Thermo Fisher Scientific).

Gene expression was quantified by qRT-PCR using PowerUp™ SYBR™ Green Master Mix and gene-targeting primers (500 nM in reaction; IDT) on a QuantStudio™ 7 Pro (Thermo Fisher Scientific). Primer sequences are shown in Supplementary Table 1. qRT-PCR data were analysed using the $\Delta\Delta C_t$ method, which compares mean $C_t$ values (from technical triplicates) to those of selected housekeeping genes (Ct average of *ACTB* and *GAPDH*), and then compares these values for test samples to those for control samples. Changes in expression are shown as $-\log_2$ fold changes.

*Immunoblotting*

To obtain whole cell lysates, cells were lysed in RIPA buffer (25 mM Tris-HCl (pH 7.4)/150 mM NaCl/1% (V/V) IGEPAL CA-630/0.5% (W/V) sodium deoxycholate/0.1% (W/V) SDS/1X protease and phosphatase inhibitor cocktails/10 µM DIFP; Merck) for 30 min on ice and the protein quantified using a Bradford assay (Thermo Fisher Scientific). Equal quantities of protein were loaded into a NuPAGE™ 4-12%, Bis-Tris, 1 mm protein gel and run at 120 V for 90 min in MOPS-SDS running buffer, alongside a PageRuler™ Plus Prestained Protein Ladder (Thermo Fisher Scientific). The protein was transferred onto a nitrocellulose membrane at 10 V for 1 hr in tris-glycine transfer buffer (50 mM tris/383 mM glycine/10% (V/V) methanol; Merck). After blocking in TBST (10 mM tris-HCl, pH 8.0/150 mM NaCl/1% (V/V) Tween20; Merck) with 5% (W/V) BSA (Merck) for 1 hr, membranes were incubated overnight at 4°C with antibody in a 5% (W/V) BSA/TBST solution, with dilutions based on the manufacturer's recommendations. The antibodies used were raised against NFATc2, total histone 3, ß-actin, COXIV and c-Myc (Supplementary Table 2). Subsequently, membranes were washed in TBST and incubated in TBST containing secondary antibodies with fluorescent conjugates (1:10,000 dilution; LI-COR) for 1 hr at RT. Membranes were visualised using a LI-COR® Odyssey Imager and analysed using ImageStudio software. Re-probing of membranes was conducted by stripping using ReBlot Plus Strong Antibody Stripping Solution (Merck), as per the manufacturer's recommendations, and re-blocking and re-probing with the appropriate antibodies as described. The antibodies used are given in Supplementary Table 2.

*RNA-seq data analyses*

The *SPIA* R package provided access to the 'signaling pathway impact analysis' (SPIA) platform, which detects enrichment of known pathways based on both the presence of genes in the data, and the strength of their deregulation relative to their position in the pathway. Other databases, such as KEGG and PANTHER were also used to establish enrichment of known pathways in the data, using their web tools as appropriate.

Scatterplots (including volcano plots) were drawn using *ggplot2* and Venn diagrams with the package *VennDiagram*, both in R. The STRING database provided information on protein-protein interactions, based on known interactions, co-expression, co-occurrence in the literature and other means of prediction.

*ChIP-seq analyses*

The *ChIPseeker* package was used in Galaxy to generate plots showing the distribution of peaks across the genome. This package also annotated peaks with Ensembl gene IDs and gene loci information. Genes were further annotated manually using the Ensembl database.

Functional enrichment in the genes containing peaks was conducted on GSEA using a pre-ranked analysis. Ensembl gene IDs were uploaded with their associated (peak) enrichment scores, which were used to rank genes. Standard settings were used and inclusion of gene sets in the range of 5-500 genes.

### Supplementary Tables

| Primer Target | Forward Primer Sequence | Reverse Primer Sequence |
|---|---|---|
| *NFATC2* | ACCCTTGGAGCCCAAAAACA | CTTTCCGCAGCTCAATGTCG |
| *GAPDH* | GTCAACGGATTTGGTCGTATTG | TGTAGTTGAGGTCAATGAAGGG |
| *ACTB* | CACAGAGCCTCGCCTTT | GCGGCGATATCATCATC |
| *MYC* | CAAGAGGCGAACACACAACG | CAACTCCGGGATCTGGTCAC |
| *CCNA2* | TGGCGGTACTGAAGTCCGG | CAAGGAGGAACGGTGACATGC |
| *CCNB1* | CAGCTCTTGGGGACATTGGTAAC | ACTGGCACCAGCATAGGTACC |
| *CCND1* | GATCAAGTGTGACCCGGACTG | CCTTGGGGTCCATGTTCTGC |
| *CCND2* | ACCAACACAGACGTGGATTGT | CTCCGACTTGGATCCGTCAC |
| *CCND3* | CCTCCTACTTCCAGTGCGTG | AGGCCAGGAAATCATGTGCA |
| *CCNE1* | CAACGTGCAAGCCTCGGA | AAAGTGCTGATCCCTTAAGTATGTC |
| *CCNE2* | ATCCTTCACCTTTGCCTGATTT | CCTCATCTGTGGTTCCAAGTCA |

**Supplementary Table 1. Primer sequences used in the study.**
Shown are the forward and reverse primer sequences used for qRT-PCR in this study, per target gene.

| Target | Species | Clone | Supplier | Catalogue ID |
|---|---|---|---|---|
| NFATc2 | Rabbit | D43B1 | CST | 5861 |
| Histone 3 | Mouse | 1B1B2 | CST | 14269 |
| B-actin | Mouse | 8H10D10 | CST | 3700 |
| COX IV | Rabbit | 3E11 | CST | 4850 |
| c-Myc | Rabbit | Y69 | Abcam | ab32072 |

**Supplementary Table 2. Western blotting antibodies used in the study.**
Each of the Western blotting antibodies used in this study is shown with the information regarding species isotype, clone, supplier and catalogue ID.

| Pathway | sh*NFATC2*-143 | | sh*NFATC2*-146 | |
|---|---|---|---|---|
| | NES | FDR q value | NES | FDR q value |
| HALLMARK_MYC_TARGETS_V1 | 2.69 | <0.001 | 2.68 | <0.001 |
| HALLMARK_MYC_TARGETS_V2 | 2.17 | <0.001 | 2.13 | 0.001 |
| HALLMARK_OXIDATIVE_PHOSPHORYLATION | 2.11 | <0.001 | 2.49 | <0.001 |
| HALLMARK_DNA_REPAIR | 1.67 | 0.010 | 1.94 | <0.001 |

**Supplementary Table 3. Five GSEA pathways were enriched in the data for sh*NFATC2* KD in THP-1 cells.** Following RNA-seq in THP-1 cells transduced with sh*NFATC2*-1, sh*NFATC2*-2, or NTC, GSEA was used to identify enriched pathways within the perturbed genes. Shown are the GSEA pathways for which the q-value was <0.05 for THP-1 both sh*NFATC2*-1 vs. NTC and sh*NFATC2*-2 vs. NTC, alongside the associated statistics: NES = normalised enrichment statistics; FDR = false discovery rate.

### Supplementary figures

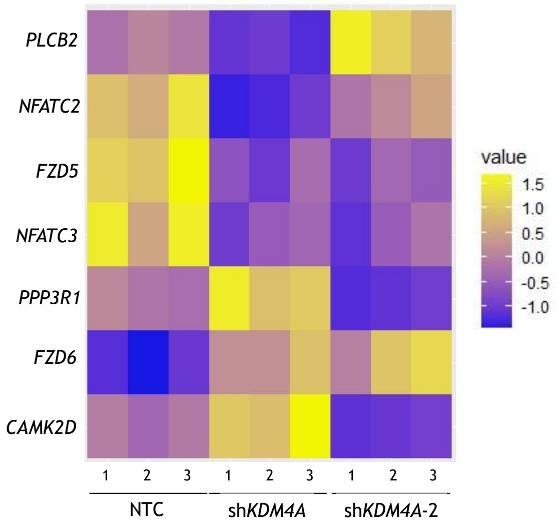


**Supplementary Figure 1. KDM4A transcriptionally regulates *NFATC2* in THP-1 cells.** As an expansion of Figure 1, THP-1 cells were transduced with shRNA constructs NTC, sh*KDM4A* or sh*KDM4A*-2. RNA-seq data were generated from cells harvested 48 hr post-puromycin selection (n=3 biological replicates). RNA-seq data from transduced THP-1 cells were analysed using the SPIA pathway enrichment tool. Shown is the expression of genes (from the topmost deregulated 'Wnt signaling pathway' from SPIA: see Figure 1A), which had $p_{adj}$<0.05 in the RNA-seq dataset for both NTC vs sh*KDM4A* and NTC vs sh*KDM4A*-2, shown as a heatmap, given as z-scaled FPKM.

---

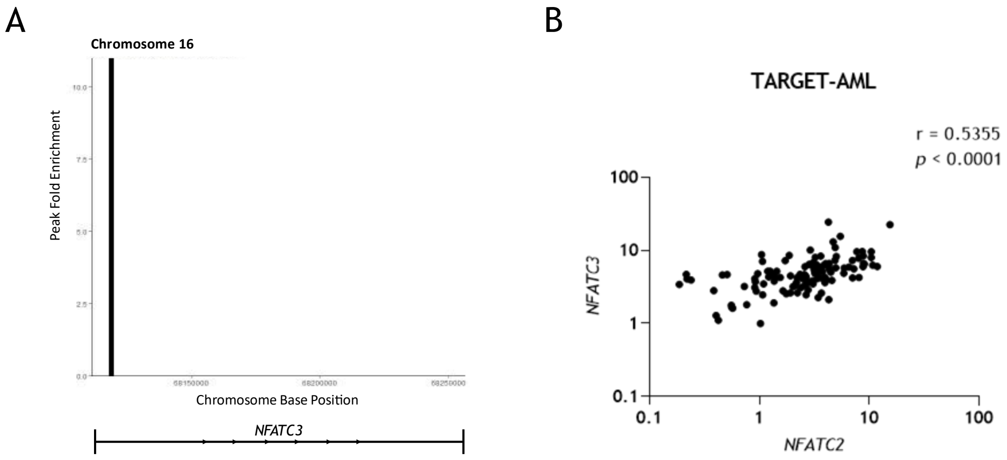


**Supplementary Figure 2. KDM4A is bound to *NFATC2* and *NFATC3* in THP-1 cells.**
**(A)** As in Figure 1, DNA/protein complexes were immunoprecipitated from untreated THP-1 cells using an anti-KDM4A antibody, and DNA was fragmented and sequenced (n=3). Shown are the KDM4A binding peaks within the *NFATC3* gene region meeting a significance threshold q<0.1, as determined by epic2. **(B)** Data from the TARGET-AML dataset were obtained using *TCGABiolinks* in R. In patient samples from BM and for which RNA-seq data were available (n=119) were compared for *NFATC2* and *NFATC3* expression. Shown is a scatterplot of expression (in FPKM) with the Pearson's correlation coefficient and *p* value of *NFATC2*–*NFATC3* correlation.

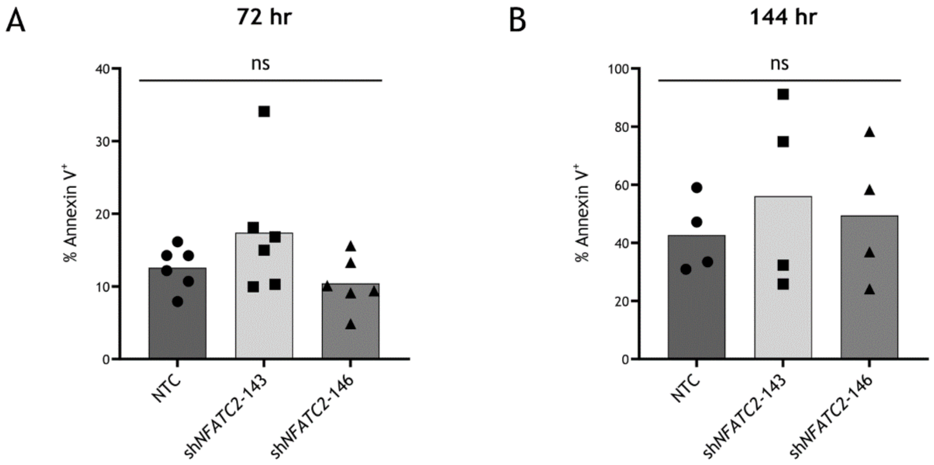


**Supplementary Figure 3. Apoptosis is not significantly increased in THP-1 cells after *NFATC2* KD.** THP-1 cells transduced with either NTC or sh*NFATC2* (-1 or -2) were stained with annexin-FITC/APC and DAPI, at either 72 hr **(A)** or 144 hr **(B)** post-puromycin selection. Shown are the mean % values of Annexin⁺ cells in each shRNA transduction group (n=3), with the associated one-way ANOVA result for an across-group comparison (ns = not significant; p>0.05).

---

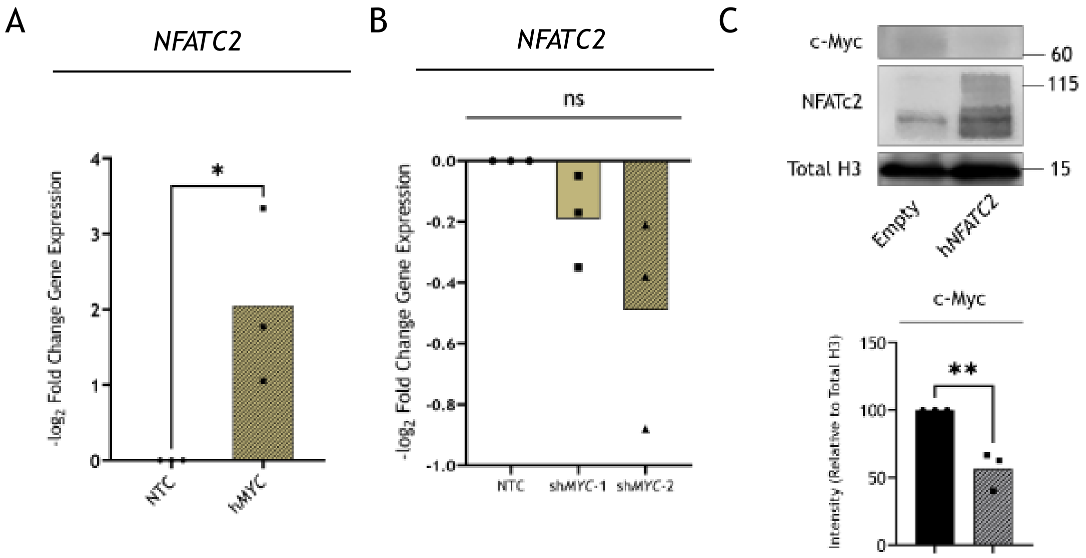


**Supplementary Figure 4. *NFATC2* OE in THP-1 cells leads to downregulation of c-Myc.**

**(A)** *NFATC2* expression measured in THP-1 cells expressing either of the vectors Empty or h*MYC* using qRT-PCR, shown as mean -$\log_2$ fold changes compared to Empty (n=3). A two-sided, unpaired t-test for a difference in means was used (*p=0.039). **(B)** *NFATC2* expression measured in shRNA-transduced THP-1 cells using qRT-PCR at 48 hr post-puromycin selection. Differential expression as compared between either NTC vs. sh*MYC*-1 and NTC vs. sh*MYC*-2 shown as mean -$\log_2$ fold changes (n=3). A one-way ANOVA for an across-group difference in means was used and the overall p value is shown (ns = not significant; p>0.05). **(C)** c-Myc protein expression was measured by immunoblot in THP-1 cells expressing either of the vectors Empty, h*NFATC2*. Quantitative densitometry results are shown (n=3), expression relative to total histone 3 (H3), and the representative immunoblot is shown. A two-sided, unpaired t-test for a difference in means was used (**p=0.006).
